# LUZP1 regulates the assembly of stress fibers by promoting maturation of contractile actomyosin bundles

**DOI:** 10.1101/2023.09.08.556811

**Authors:** Liang Wang, Hoi Ying Tsang, Ziyi Yan, Sari Tojkander, Katarzyna Ciuba, Konstantin Kogan, Xiaonan Liu, Hongxia Zhao

**Author notes:** Corresponding author: Dr. Hongxia Zhao, Faculty of Biological and Environmental Sciences, University of Helsinki, 00014 Helsinki, Finland, Finland.

## Abstract

Contractile actomyosin bundles play crucial roles in various physiological processes, including cell migration, morphogenesis, and muscle contraction. The intricate assembly of actomyosin bundles involves the precise alignment and fusion of myosin II filaments, yet the underlying mechanisms and factors involved in these processes remain elusive. Our study reveals that LUZP1, a leucine zipper protein, plays a central role in orchestrating the formation of thick actomyosin bundles. Loss of LUZP1 caused abnormal cell morphogenesis, migration, and the ability to exert forces on the environment. Importantly, knockout of LUZP1 results in significant defects in the concatenation and persistent association of myosin II filaments, severely impairing the assembly of myosin II stacks. The disruption of these processes in LUZP1 knockout cells provides mechanistic insights into the defective assembly of thick ventral stress fibers and the associated cellular contractility abnormalities. Overall, these results significantly contribute to our understanding of the molecular mechanism involved in actomyosin bundle formation and highlight the essential role of LUZP1 in this process.

## Introduction

The actin cytoskeleton plays a fundamental role in numerous essential cellular processes, such as migration, morphogenesis, cytokinesis, and endocytosis. To facilitate these processes, actin filaments assemble into structures that protrude and contract, generating forces through actin polymerization against the plasma membrane and the ATP-dependent sliding of myosin II filaments along actin filaments. In animal cells, contractile actomyosin structures encompass the cytokinetic contractile ring, myofibrils in muscle cells, and stress fibers in non-muscle cells. Stress fibers comprise actin, non-muscle myosin II (NM-II), and various actin-binding proteins (Beach et al., 2017; Betapudi, 2014; Betapudi et al., 2010). Based on their protein compositions and associations with focal adhesions, stress fibers are categorized into distinct subtypes, including dorsal stress fibers, transverse arcs, ventral stress fibers, and the perinuclear actin cap or actin cap (Tojkander et al., 2012). Among these, transverse arcs, ventral stress fibers, and the actin cap are contractile actomyosin bundles containing the motor protein NM-II. NM-II plays a pivotal role in diverse cellular processes, including cell motility, contractility, and maintenance of cell shape (Somlyo and Somlyo, 2000; Vicente-Manzanares et al., 2009). It generates force through the ATP-dependent interaction between its motor domain and actin filaments. Alongside its contractile functions, NM-II also cross-links and stabilizes actin by directly binding to actin filaments (Siddique et al., 2005). NM-II is a bipolar, contractile protein composed of two myosin heavy chains (MHCs), two regulatory myosin light chains (RLCs), and two essential light chains (ELCs). Each MHC consists of an N-terminal globular motor domain responsible for driving filament sliding, and a C-terminal tail/rod region that associates with the other MHC (Landsverk and Epstein, 2005). The contractility of stress fibers is meticulously regulated through the phosphorylation of the myosin light chain. This phosphorylation event enhances the assembly of myosin II filaments and stimulates the actin-activated ATPase activity of the myosin motor domain (Somlyo and Somlyo, 2000; Vicente-Manzanares et al., 2009).

Previous studies have demonstrated that thick actomyosin bundles form through the lateral fusion of transverse arcs, with a preference for fusion at the intersections with dorsal stress fibers. This fusion process is accompanied by the alignment of distal focal adhesions (Hotulainen and Lappalainen, 2006; Tojkander et al., 2015). The development of larger NM-II macromolecular assemblies, known as NM-II stacks, occurs through the expansion of initial NM-II filaments and stacks, as well as the concatenation of NM-II filaments. The development of larger NM-II macromolecular assemblies, known as NM-II stacks, occurs through the expansion of initial NM-II filaments and stacks, as well as the concatenation of NM-II filaments (Beach et al., 2017; Hu et al., 2017). These processes depend on NM-II motor activity and actin filament assembly, both of which are crucial for organizing arcs into ventral stress fibers. While the mechanisms governing the assembly of protrusive actin filament networks are relatively well-established, the intricate process of forming NM-II-containing contractile filament structures is not fully understood (Lehtimäki et al., 2017; Ono, 2010; Sophie Mokas et al., 2009; Vicente-Manzanares et al., 2007).

Leucine zipper protein 1 (LUZP1) was initially recognized for its significance in embryonic development, as the knockout of LUZP1 in mice resulted in severe neural tube closure defects during brain development, ultimately leading to perinatal death (Gonçalves et al., 2020; Hsu et al., 2008). This underscores the critical importance of LUZP1 for survival. LUZP1 has been observed to localize to various cellular locations, including the centrosome, the basal body of primary cilia, the midbody, actin filaments, and cellular junctions (Bozal-Basterra et al., 2020; Gonçalves et al., 2019, 2020; Wang and Nakamura, 2019a; Wang and Nakamura, 2019a, 2019b; Yano et al., 2021). Previous studies have shown that LUZP1 regulates the formation of primary cilia (Bozal-Basterra et al., 2020; Gonçalves et al., 2019, 2020; Wang and Nakamura, 2019a). Additionally, LUZP1 has been identified as a microtubule-associated protein found at circumferential rings within the tight junctions of epithelial cells, where it is important for apical constriction (Yano et al., 2021). Recently, LUZP1 has been implicated in controlling cell division, migration and invasion through the regulation of the actin cytoskeleton (Bozal-Basterra et al., 2021). However, the molecular details and mechanisms by which LUZP1 contributes to stress fiber assembly, maintenance, and contractility in non-muscle cells remain to be elucidated.

In this study, we demonstrate that LUZP1 plays an essential role in the maturation of thick contractile stress fibers by specifically localizing to the neck region of myosin II. Depletion of LUZP1 results in significant defects in the concatenation and persistent association of myosin II filaments, leading to a severe impairment in the assembly of myosin II stacks. Consequently, the absence of LUZP1 disrupts the maturation of actomyosin bundles from their precursors, with direct implications for cell migration and the ability of cells to exert forces on their environment. Overall, our findings underscores LUZP1’s significance as a key regulator of stress fiber dynamics, establishing its connection to fundamental cellular processes. By orchestrating the assembly of myosin II filaments and filament stacks within stress fibers, LUZP1 directly influences their structural integrity and functionality.

## Results

### LUZP1 Is Component of Contractile Actin Stress Fibers

LUZP1 has been experimentally identified to have two splice variants (1076 aa and 1026 aa) that share identical sequences between residues 1-1024 aa, but have distinct C-terminal regions (Gonçalves et al., 2020; Wang and Nakamura, 2019a)(Figure 1A). To assess the expression levels of these LUZP1 isoforms in U2OS cells, real-time quantitative PCR (RT-qPCR) analysis was performed. The results showed that both isoforms are expressed in U2OS cells, with the longer isoform, LUZP1-1076 aa, being approximately six times more abundant at the mRNA level (Figure 1B). To examine the actin-subtype localization of LUZP1, immunofluorescence microscopy was employed using an anti-LUZP1 antibody generated against residues 334-427 of LUZP1, which can recognize both isoforms. The results revealed that LUZP1 localized in three subtypes of actin stress fibers, including dorsal stress fibers, transverse arcs, and ventral stress fibers (Figure 1C). Particularly, LUZP1 was enriched in the thick actomyosin bundles, suggesting its specific role in the formation and organization of these contractile structures. To corroborate the stress fiber localization of LUZP1, we expressed GFP-tagged LUZP1 in U2OS cells. In concordance with the antibody staining, GFP-tagged LUZP1 prominently resided within stress fibers, notably concentrated in the thick contractile actomyosin bundles (Figures 1D and S1). Furthermore, the expression of LUZP1-GFP led to a significant increase in the number of thick stress fibers, strongly indicating that LUZP1 actively contributes to the formation of these thick actomyosin bundles (Figure 1D). Additionally, a single bright punctum besides the stress fiber localization was observed (Figures S1 and 1C). The sporadic nuclear localization shown in the LUZP1 antibody staining was absent in LUZP1-GFP-expressing cells (see Figures S1 and 1C), which hints at the possibility of nonspecific antibody binding accounting for the sporadic nuclear staining.

**Figure 1.**
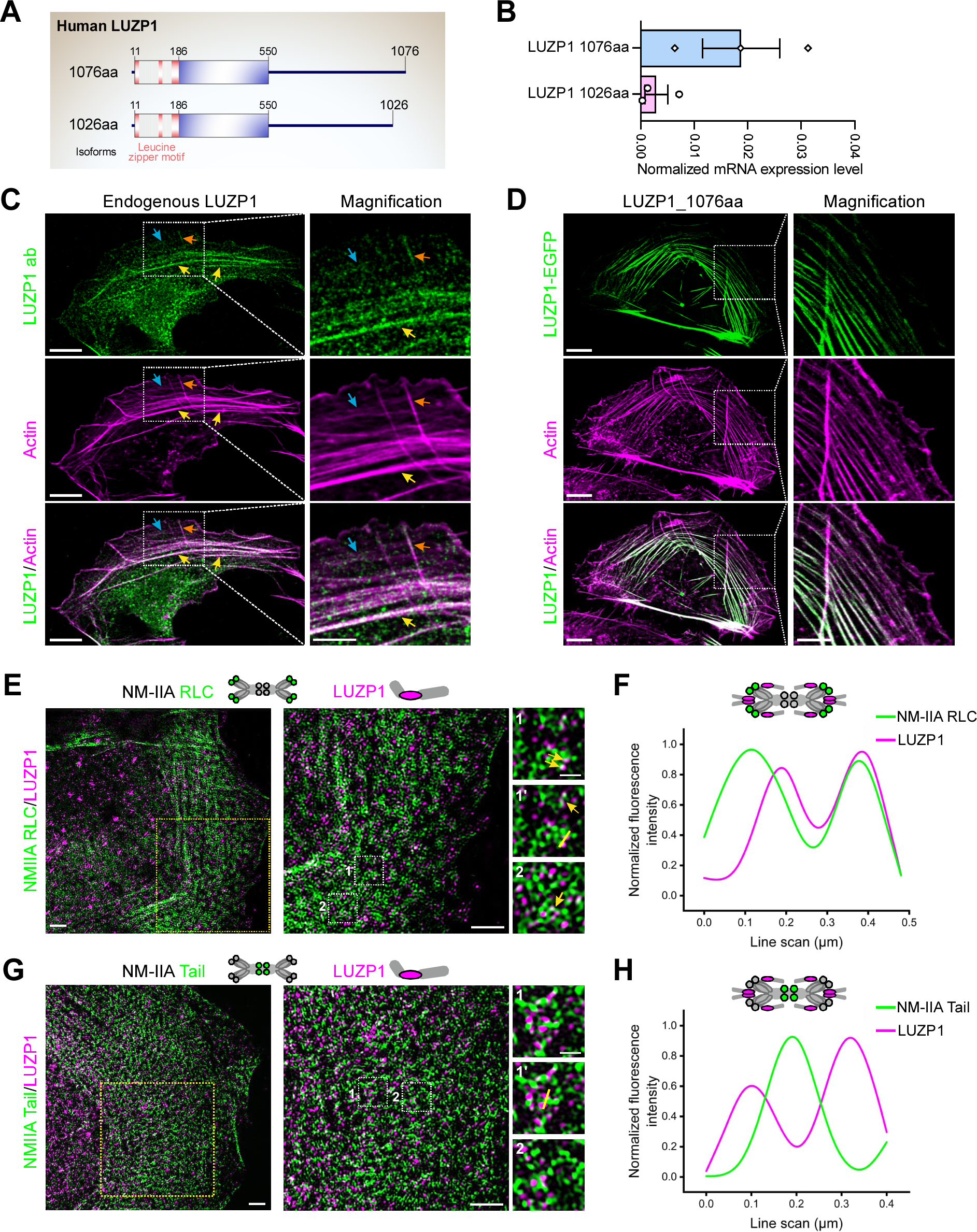
LUZP1 localizes in close proximity to the neck region of myosin II. (**A**) Illustration of domain structures of two LUZP1 isoforms in U2OS cells. Two isoforms are identical between 1-1024 aa containing three leucine zipper motifs at the N-terminus. (**B**) Real-time quantitative PCR (RT-qPCR) analysis of the relative transcription levels of LUZP1 isoforms. (**C**) Cellular localization of the endogenous LUZP1 by immunofluorescence microscopy analysis. Scale bar: 10 µm (main), and 5 µm (insets). (**D**) Cellular localization of GFP tagged LUZP1 by immunofluorescence microscopy analysis. F-actin was visualized by fluorescent phalloidin. Actin stress fibers including the transverse arcs, dorsal and ventral stress fibers were indicated by blue, orange, and yellow arrows. Scale bar: 10 µm (main), and 5 µm (insets). (**E**) Representative 3D-SIM images of a U2OS cell expressing mApple-NM-IIA RLC (green) and stained with LUZP1-specific antibody (magenta). Boxes with yellow and white dotted lines indicate the magnified regions. The magnified image 1’ shows an NM-IIA filament/LUZP1 that was used for line plot analysis. **(F)** Line plot analysis of the co-localization of LUZP1 and NM-IIA RLC. (**G**) Representative 3D-SIM image of a U2OS cell expressing mEmerald-NM-IIA tail (green) and stained with LUZP1-specific antibody (magenta). Boxes with yellow and white dotted lines indicate the magnified regions. The magnified image 1’ shows an NM-IIA filament/LUZP1 that was used for line plot analysis. Scale bars for panels F and G: 2 µm for the left and middle panels, 1 µm for the right panel. **(H)** Line plot analysis of the co-localization of LUZP1 and NM-IIA tail.

LUZP1 is a large protein (120 kD) containing three leucine zipper motifs at its N-terminus (Figure S2A). To investigate the contributions of different regions of LUZP1 to its cellular localization, various GFP-tagged deletion constructs of LUZP1 were generated and expressed in U2OS cells. The results showed that the N-terminal region of LUZP1, including the leucine zipper motifs (LUZP_1-252), did not localize to stress fibers (Figure S2B). However, a longer N-terminal extension (LUZP_1-550) displayed stress fiber localization similar to that of the full-length LUZP1, primarily residing in thick actomyosin bundles (Figure S2B). On the other hand, the middle region (LUZP1_252-550) and C-terminal extension (LUZP_550-1076), lacking the N-terminal leucine zipper motif, localized to thin stress fibers (Figure S2B). These data suggest that localization of LUZP1 to the thick actomysin bundles requires both the leucine zipper and the actin binding motifs. Taken together, these findings provide experimental evidence that LUZP1 is a stress fiber-associated protein specifically favoring the formation of thick actomyosin bundles. Its expression and localization in actin stress fibers, especially in thick contractile actomyosin bundles, suggest its involvement in modulating the organization and architecture of the actin cytoskeleton.

### LUZP1 Localizes in Close Proximity to the Neck Region of the Non-muscle Myosin II

To unravel the precise association of LUZP1 with stress fiber components, we employed super-resolution structured-illumination microscopy (SIM). The results corroborated the data from conventional fluorescence microscopy, showing that LUZP1 is localized to F-actin-rich stress fibers, with the highest abundance observed in mature ventral stress fibers and thick arc bundles. While LUZP1 did not exhibit co-localization with the actin crosslinking protein α-actinin and instead displayed periodic staining with α-actinin (Figure S3A), super-resolution microscopy unveiled a close localization of LUZP1 with the regulatory light chain (RLC) of NM-II (Figures 1E, 1F, and S3B). This localization pattern indicates that LUZP1 is positioned in close proximity to the neck region of NM-II in stress fibers. The line plot displaying the co-localization of LUZP1 and NM-IIA RLC demonstrated the close association between LUZP1 and NM-II in stress fibers (Figures 1E and 1F). In addition to its localization within thick actomyosin bundles, the close proximity observed between LUZP1 and NM-II strongly suggests that LUZP1 plays a crucial role in the assembly, organization, and contractility of stress fibers. This inference gains further support from the absence of co-localization between LUZP1 and the NM-IIA tail, as visualized by the mEmerald-NM-IIA tail construct (Figures 1G and 1H), indicating the specificity of LUZP1’s association with the neck region of NM-II. Taken together, the super-resolution microscopy data provides valuable insights into the precise localization of LUZP1 within stress fibers and its potential role in the regulation of NM-II.

### LUZP1 Regulates Maturation of Actomyosin Bundles

To investigate the cellular role of LUZP1, we initially conducted loss-of-function experiments using small interfering RNA (siRNA) silencing. Validation through real-time qPCR and western blot analyses confirmed the efficient downregulation of both LUZP1 isoforms, evident at both the transcriptional and translational levels (Figures 2A, S4A, and S4B). Additionally, we generated LUZP1 knockout U2OS cells via CRISPR-Cas9 technology. A subsequent western blot analysis validated the absence of detectable LUZP1 protein in these knockout cells (Figure 2A). To further confirm the LUZP1 knockout, we conducted sequencing analysis, revealing a deletion of 2079 nucleotides from the LUZP1 genome DNA, which corresponds to amino acids 1-664 aa (Figures S4C and S4D). These data collectively demonstrate the successful knockout of LUZP1. Immunofluorescence microscopy showed that both the LUZP1 knockout and knockdown cells lacked LUZP1 staining in the stress fibers (Figure S5A and S5B). The sporadic staining observed in the nucleus can be attributed to the unspecific binding of the antibody, as evidenced by the non-nuclear localization of LUZP1-GFP (Figures S1B, S5A, and S5B).

**Figure 2.**
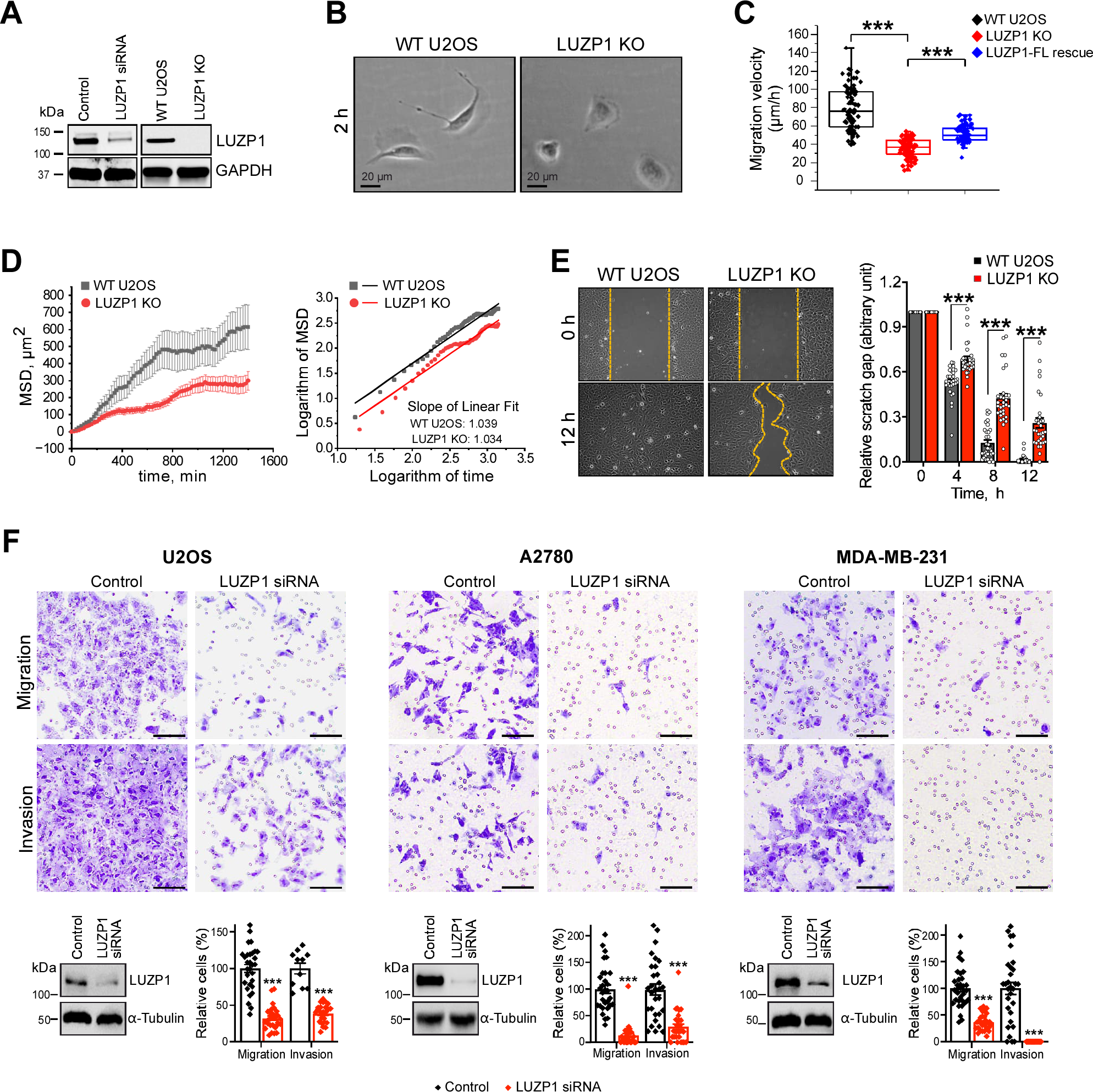
Loss of LUZP1 caused defects in cell migration. (**A**) Western blot analysis of endogenous LUZP1 protein levels after 72h treatment with control or LUZP1 target-specific siRNA in U2OS cells, as well as the LUZP1 protein levels in wild-type U2OS and LUZP1 knockout cells. GAPDH was probed for equal sample loading. (**B**) Representative bright field images of wild-type and LUZP1 knockout cells 2h after the cells were plated on a fibronectin-coated surface. Scale bars, 20 µm. (**C**) Random migration velocities of wild-type and LUZP1 knockout cells. Quantification is based on manual tracking of the displacement of nuclei within 12 h with 20 min interval. n = 82 for wild-type and n = 94 for LUZP1 knockout cells. ***P≤0.001, Student’s *t* test. (**D**) Mean square displacement (MSD) analysis of cell migration. (**E**) Wound healing assay for wild-type and LUZP1 knockout cells. Representative bright field images of start (0 h) and end (12 h) time point were shown in the upper panel. The statistical analysis of the gap distance in relation to the starting point is presented in the right panel. n = 31 for both wild-type and LUZP1 knockout cells. Error bars indicate ±SEM. ***. P≤0.001, Student’s *t* test. (**F**) The effects of LUZP1 silencing by siRNA on cell migration and invasion in different cell lines.

Morphologically, LUZP1 knockout cells appeared slightly smaller and displayed a more round shape compared to wild-type U2OS cells when plated on a fibronectin-coated cover glass and incubated for 2 hours before imaging (Figure 2B). Strikingly, the LUZP1 knockout cells exhibited significant migratory defects. The random migration velocity and wound healing ability of LUZP1-knockout cells were significantly reduced compared to wild-type cells, as demonstrated in Figures 2C-2E. To further analyze cell migration patterns, Mean Squared Displacement (MSD) analysis was performed using the cell trajectories obtained from time-lapse data. The MSD analysis revealed that wild-type U2OS cells explored a significantly larger area compared to LUZP1 knockout cells (Figure 2D). The log-log plot of MSD as a function of time increment showed that both wild-type and LUZP1 knockout cells exhibited random migration, with exponents α of 1.039 and 1.034, respectively (Figure 2D). The differences in MSD between wild-type and LUZP1 knockout cells indicate alterations in the migratory behavior and dynamics of LUZP1-deficient cells. To examine whether the effects of LUZP1 depletion on cell migration are cell-specific, siRNA treatment was performed in different cell lines, including U2OS, A2780, and MAD-MB-231 cells (Figure 2F). The results revealed that siRNA-mediated knockdown of LUZP1 consistently led to impaired cell migration and invasion ability in all three cell lines (Figure 2F). These findings indicate that the role of LUZP1 in regulating cell migration and invasion is a general mechanism across the different cell lines we tested.

Importantly, in LUZP1-knockout cells, the transverse arcs displayed significant defects in assembling into thick, contractile actomyosin bundles (Figure 3A). These cells showed an increased number of thin transverse arcs and a diminished number of thick actomyosin bundles, indicating a disruption in the assembly of contractile stress fibers. To analyze the stress fiber phenotype more precisely, cells were plated on crossbow-shaped fibronectin micro-patterns, which allowed for a characteristic organization of the stress fiber network (Tojkander et al., 2015). In line with the findings in cells plated on normal substrates, LUZP1 knockout cells on micro-patterns exhibited a higher number of thin arcs and a reduced number of thick actomyosin bundles compared to control cells (Figure 3B). Corresponding stress fiber phenotypes were similarly observed in U2OS cells treated with LUZP1 siRNA (Figures 3A and 3B), providing additional confirmation of LUZP1’s role in facilitating the assembly of thick actomyosin bundles.

**Figure 3.**
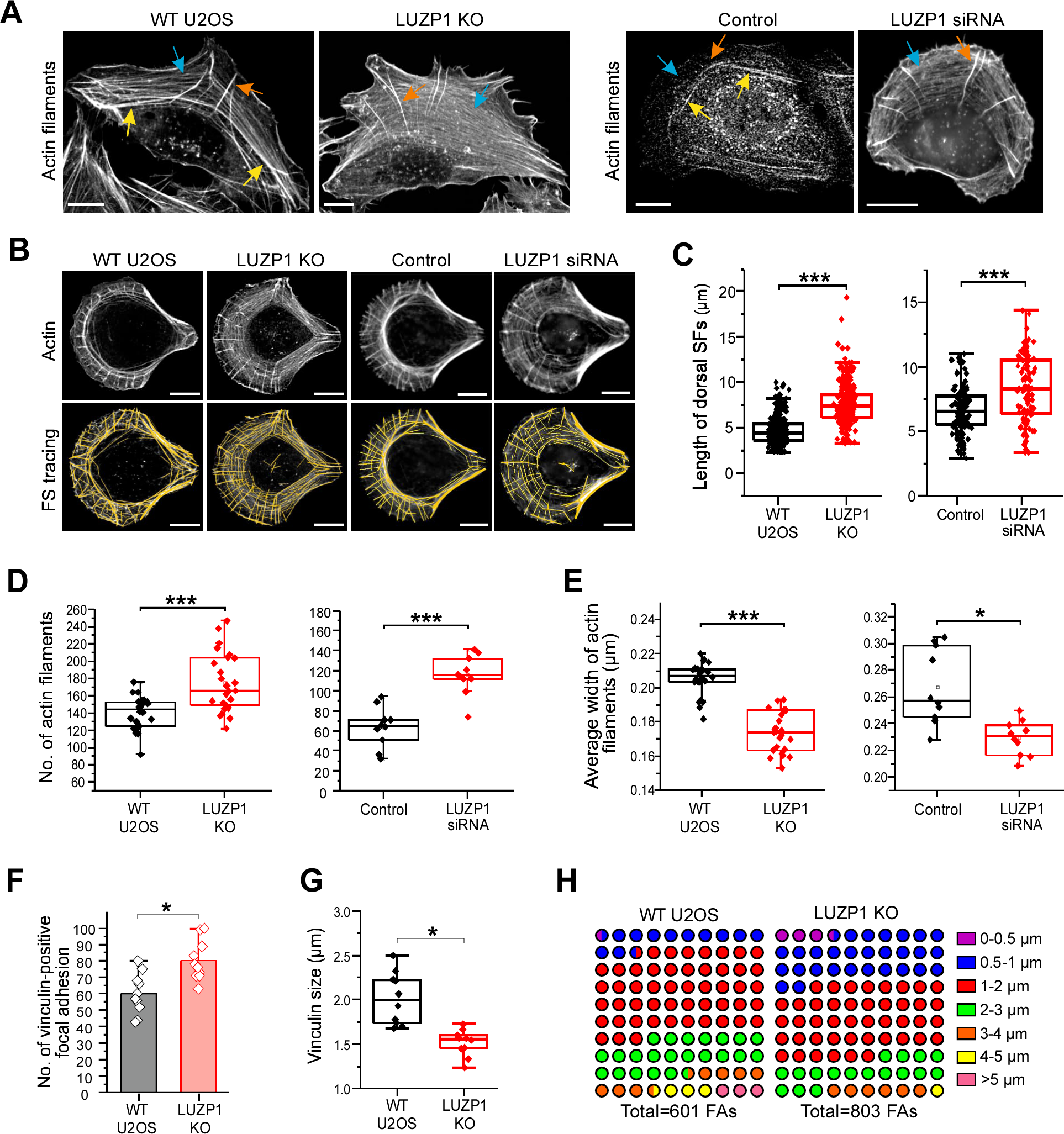
Loss of LUZP1 led to defects in the maturation of stress fibers. (**A**) Representative images of actin filaments visualized by fluorescent phalloidin in wild-type, LUZP1 knockout and knockdown cells. After treatment with siRNA for 72 h, cells were re-plated on fibronectin coated cover glass and further cultured for 2 h before fixation. Examples of dorsal stress fibers, transverse arcs, and ventral stress fibers are indicated by orange, blue, and yellow arrows, respectively. Scale bars, 10 µm. (**B**) Representative images of actin filaments visualized by fluorescent phalloidin in wild-type, LUZP1 knockout and knockdown cells cultured on crossbow shaped micro-patterns coated with fibronectin (upper panel). The corresponding tracing images of actin filaments using the software FilamentSensor_0.2.2 were shown in the lower panel. Scale bars, 10 µm. (**C**) Quantification of the dorsal stress fiber length in wild-type, LUZP1 knockout and knockdown cells. n=310 dorsal SFs from 24 cells (wild-type), n=316 dorsal SFs from 26 cells (LUZP1 knockout), n=124 dorsal SFs from 7 cells (Control scramble) and n=96 dorsal SFs from 9 cells (LUZP1 knockdown). ***, P≤0.001, Student’s *t* test. (**D&E**) Quantification of the number (D) and average width (E) of traced actin filaments in wild-type, LUZP1 knockout and knockdown cells. n=24 for wild-type, n=26 for LUZP1 knockout cells, n=124 for control and n=96 for LUZP1 knockdown cells. ***, P≤0.001, Student’s *t* test. (**F&G**) Quantification of the average number and size of vinculin-positive focal adhesions (FAs) in wild-type and LUZP1 knockout cells. n = 10 cells for both wild-type and LUZP1 knockout cells. *, P≤0.05, Student’s *t* test. (**H**) The 10 x 10 dot plot shows the ratio of each group of vinculin-positive FAs in wild-type and LUZP1 knockout cells. n = 601 FAs from 10 wild-type cells and n = 803 FAs from 10 LUZP1 knockout cells.

Furthermore, LUZP1-deficient cells displayed notably elongated dorsal stress fibers, which is a characteristic phenotype for U2OS cells lacking thick ventral stress fibers (Figures 3A-3C) (Tojkander et al., 2015). Quantification analysis of this stress fiber phenotype in LUZP1-deficient cells using FilamentSensor software (Eltzner et al., 2015) revealed a significant increase in the number of actin filaments, coupled with a reduced average width of these filaments (Figures 3B, 3D and 3E). This observation is consistent with augmented presence of thin stress fibers characteristic of LUZP1 knockout cells. Furthermore, quantification analysis showed that LUZP1 knockout cells exhibited an elevated number of focal adhesions, while the average size of vinculin-positive focal adhesions decreased (Figures 3F-3H and S5C). These findings underscore the disruption of stress fiber assembly and dynamics in the absence of LUZP1. Indeed, in wild-type U2OS cells, transverse arcs initially appeared near the leading edge and fused together as they progressed toward the cell center (Figures 4A, Videos S1 and S2). However, in LUZP1 knockout cells, the transverse arcs failed to effectively fuse as they moved toward the cell center. This fusion impairment was accompanied by a significant reduction in the centripetal flow rates of the arcs (Figure 4B). Moreover, the actin retrograde flow rate in the lamellipodia was significantly diminished in LUZP1 knockout cells (Figure 4C). Significantly, both LUZP1 knockdown and knockout cells demonstrated notably weakened contractile forces when compared to control cells (Figures 4D and 4E). This observation suggests that the impaired fusion of transverse arcs in LUZP1-deficient cells directly contributes to the reduction in their ability to generate contractile forces and exert mechanical effects on the surrounding environment. These findings underscore the pivotal role of LUZP1 in facilitating the fusion of transverse arcs and subsequently driving the generation of contractile forces within cells.

**Figure 4.**
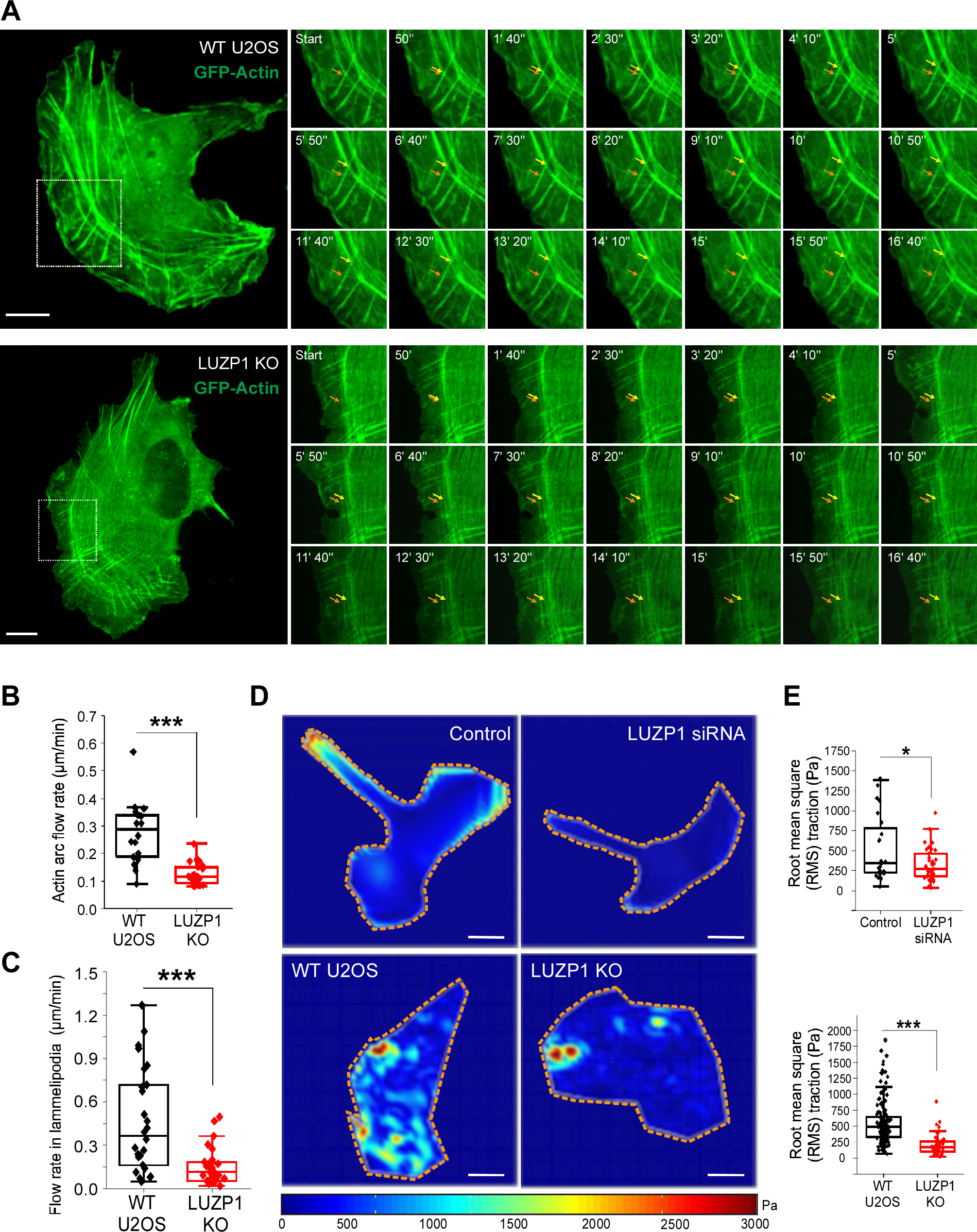
LUZP1 knockout results in disruption of transverse arc fusion during centripetal flow and decreased contractile force. (**A**) Representative examples of transverse arc flow visualized in wild type and LUZP1 knockout U2OS cells expressing GFP-actin. The orange arrows indicate the positions of the observed transverse arcs in the beginning of the movies (start), and yellow arrows indicate the positions of the same arcs in subsequent time-lapse images. Scale bar: 10 µm. (**B**) The average centripetal flow rates of transverse arcs in wild-type (n = 18) and LUZP1 knockout (n = 25) cells. ***, P≤0.001, Student’s *t* test. (**C**) The actin retrograde flow rate in the lamellipodia in wild-type (n = 26) and LUZP1 knockout (n = 31) cells. ***, P≤0.001, unpaired two-tailed *t* test. (**D**) Force maps of representative wild-type, LUZP1 knockout cells, as well as control siRNA and LUZP1 siRNA treated U2OS cells, grown on 25-kPa dishes with fluorescent nanobeads. Scale bar: 20 µm (**E**) Quantification of traction forces (root mean square traction) in wild-type (n = 24), LUZP1 knockout cells (n = 43), control (n=148) and LUZP1 knockdown cells (n = 54). *, P≤0.05; **, P≤0.001, Student’s *t* test.

Furthermore, LUZP1 knockout cells exhibited other distinct phenotypic characteristics, including a broader lamella, increased cell height, and the repositioning of nuclei towards the rear of the cell (Figures 5A and 5B). These features are commonly associated with deficiencies in stress fiber assembly and contractility. Notably, the phenotypic stress fiber alterations in LUZP1 knockout cells could be restored by the introduction of full-length LUZP1-GFP into these cells (Figure 5C and 5D). LUZP1-knockout cells expressing full-length LUZP1-GFP exhibited normal stress fiber architecture as the wild type cells (Figure 5D), supporting the notion that the observed phenotypes in LUZP1-knockout cells can be directly attributed to the deficiency of LUZP1. Furthermore, the longer N-terminal extension of LUZP1 (LUZP1_1-550) was able to efficiently rescue the assembly of thick actomyosin bundles in LUZP1 knockout cells, similarly to the rescue effect of the full-length LUZP1 (Figure 5D). In contrast, the other mutants (LUZP_1-252, LUZP1_252-550, and LUZP1_550-1076) failed to rescue the assembly of thick actomyosin bundles (Figure 5D). These findings indicate that the N-terminal extension (LUZP1_1-550) is critical for the assembly of thick contractile actomyosin bundles. In contrast, the C-terminal region of LUZP1 plays a minor role in this regard.

**Figure 5.**
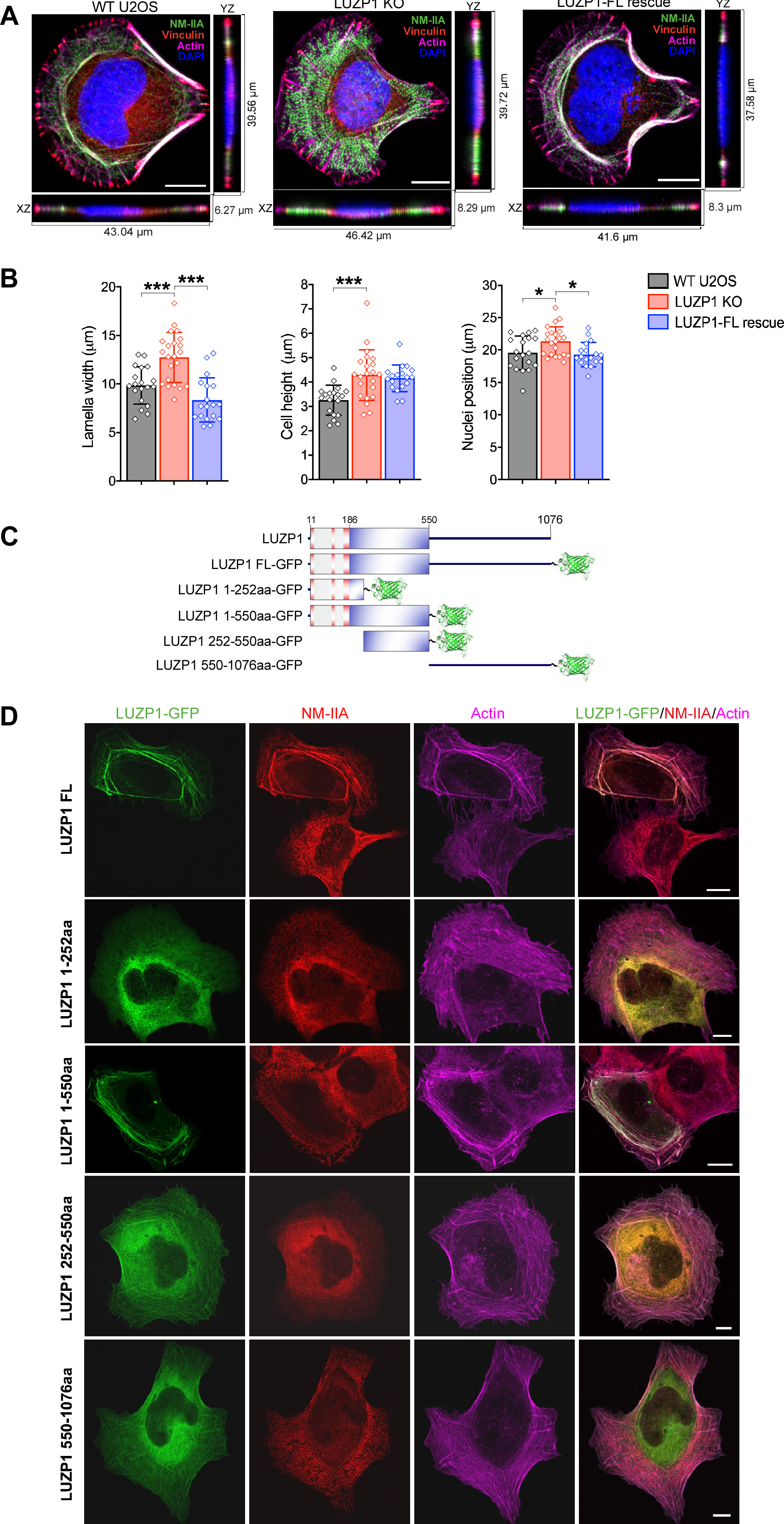
Defects in the maturation of stress fibers by loss of LUZP1 can be rescued by expressing LUZP1 in LUZP1 knockout cells. (**A**) Maximum z-projections and side views of wild type and LUZP1 knockout cells, as well as LUZP1 knockout cells expressing the full-length LUZP1 (LUZP1-FL) which were plated on the fibronectin coated crossbow shaped micropatterns. Cells were visualized for NMIIA, Vinculin, F-actin and DAPI. Scale bar: 10 µm. (**B**) Quantification of lamella width, cell height and distance of nuclei from the leading edge in wild-type, LUZP1 knockout and LUZP1-Full length (FL) rescue cells. n = 19 for wild-type, 22 for LUZP1 knockout cells and 18 for LUZP1-FL rescue cells. *, P≤0.05; **, P≤0.01; ***, P≤0.001, Student’s *t* test. (**C**) Schematic diagram of domain structures of GFP-tagged LUZP1 constructs. (**D**) Representative images for the rescue of stress fiber maturation defects in LUZP1 knockout cells by expressing wild type and mutant proteins of LUZP1.

### LUZP1 Regulates the Phosphorylation of Myosin II and the Targeting Subunit of the Myosin Phosphatase MYPT1

Cell migration and the ability to exert forces on the environment rely on the presence of contractile actin bundles containing myosin. Given LUZP1’s co-localization with myosin, we aimed to uncover the underlying causes of the disruptions in stress fiber network organization and contractility observed upon LUZP1 loss. For this purpose, we initially investigated the distribution of NM-II in both control and LUZP1 knockout cells. Immunofluorescence microscopy showed that NM-II in LUZP1 knockout cells exhibited a scattered and punctate stress fiber localization pattern, in contrast to the distinct thick stress fiber localization seen in control cells (Figures 6A and S6A). This observation is consistent with the increased number of thin stress fibers and loss of thick actomyosin bundles in LUZP1 knockout cells. To further examine whether LUZP1 regulates the association of myosin II with stress fibers, we conducted an analysis of the abundance of myosin II incorporated into stress fibers in LUZP1-knockout cells. Immunostaining of wild-type and LUZP1-knockout cells using antibodies against NMIIA or NMIIB revealed no significant alterations in the intensity of NM-IIA or NM-IIB in stress fibers of LUZP1-depleted cells (Figures 6A and S6A). Moreover, the overall protein levels of NMIIA and NMIIB in cells were not affected by LUZP1 knockout (Figure 6B).

**Figure 6.**
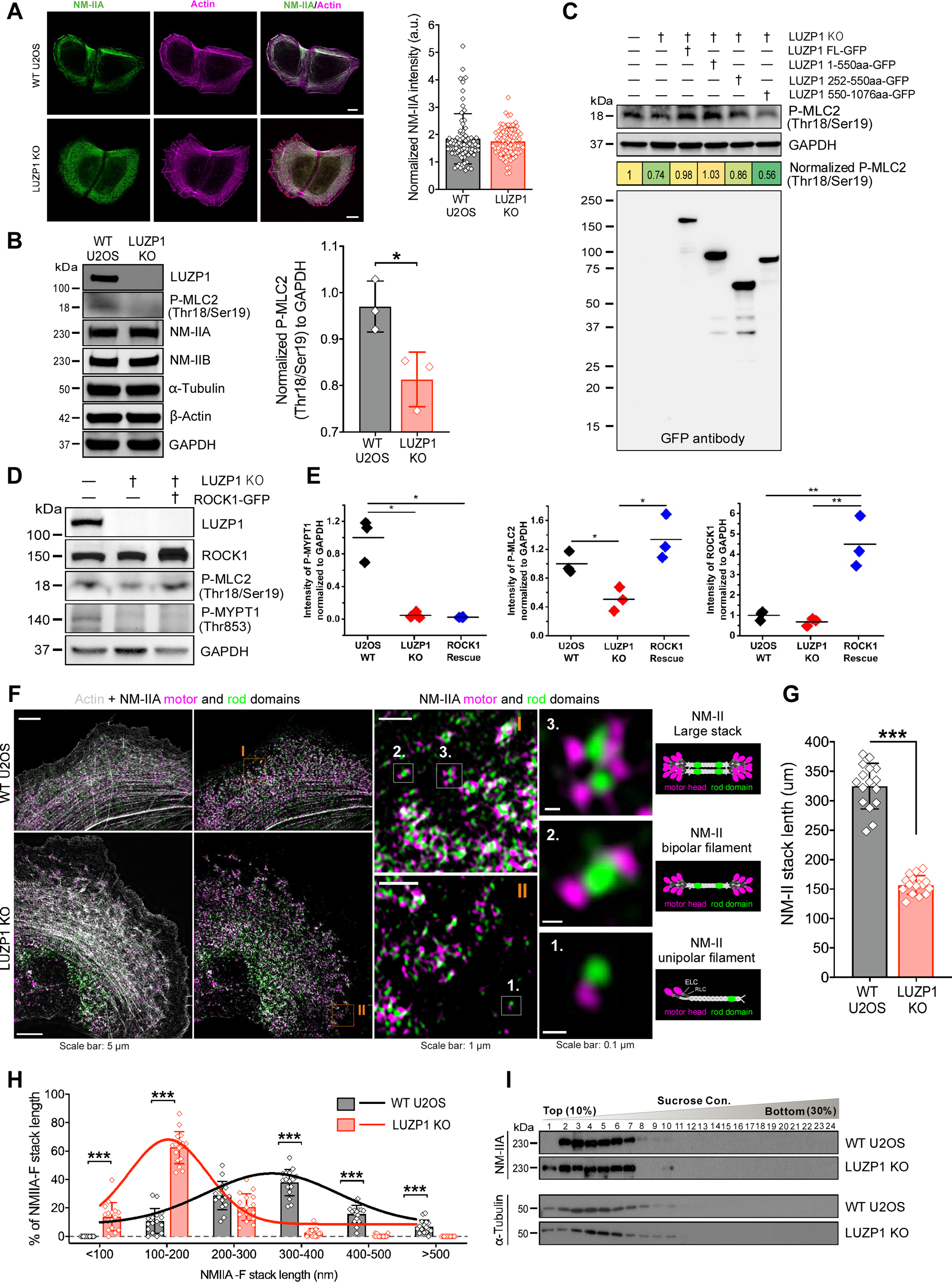
Loss of LUZP1 results in defects in the phosphorylation of the targeting subunit of the myosin phosphatase MYPT1 and the assembly of large NMII stacks. (**A**) Representative images of myosin IIA visualized by myosin II heavy chain antibody in wild-type and LUZP1 knockout cells grown on fibronectin coated cover glasses. The actin filaments are visualized by fluorescent phalloidin. Scale bars, 10 µm. The myosin intensity was normalized to the actin intensity (right panel). n = 77 for wild-type cells and n = 83 for LUZP1 knockout cells. (**B**) Western blotting analysis of endogenous myosin II (NM-IIA and NM-IIB) and double-phosphorylated regulatory light chain (P-MLC2 Thr18/Ser19) of myosin II in total cell lysates of wild type and LUZP1 knockout U2OS cells. GAPDH was probed for equal sample loading. The double phosphorylated protein levels of MLC2 were quantified by the intensity of P-MLC2 Thr18/Ser19 band normalized to the corresponding GAPDH intensity (right panel). n = 3 for both wild-type and LUZP1 knockout cells. **, P≤0.01, Student’s *t* test. (**C**) Rescue of myosin II phosphorylation by expressing LUZP1 and its truncated mutants in LUZP1 knockout cells. (**D**) Western blotting analysis of myosin RLC phosphorylation (Thr18/Ser19), phosphorylation of the myosin phosphatase targeting subunit MYPT1 (Thr853), and ROCK1 in wild type U2OS cells, LUZP1 knockout cells, and LUZP1 knockout cells expressing ROCK1-GFP. GAPDH was probed for equal sample loading. Quantification was performed by normalizing each protein intensity to the corresponding GAPDH intensity (right panel). n = 3. *, P≤0.05, **, P≤0.01, Student’s *t* test. (**E**) Quantification of myosin RLC phosphorylation (P-MLC2), phosphorylation of the myosin phosphatase targeting subunit MYPT1 (P-MYPT1), and ROCK1 in Panel D. (**F**) Representative 3D-SIM images of NMIIA stack formation in wild-type and LUZP1 knockout cells. Actin filaments were visualized by fluorescent phalloidin. NMIIA motor and tail/rod domains were visualized by a mApple-MyosinIIA-C-18 construct and an antibody against NM-IIA C-terminus, respectively. Scale bar: 5 µm. Magnified images (corresponding to the orange boxes) display characteristic NM-IIA filament distributions in wild type and LUZP1 knockout cells. Scale bar: 1 µm. High-magnification images of individual filaments (corresponding to the white boxes in the magnified regions) display examples of unipolar structure (1); bipolar filament (2); and stack of bipolar filaments (3). Scale bar: 0.1 µm. (**G&H**) Quantification of the average length (µm) (G) and the distributions (H) of NMIIA stacks. n = 641 for 10 wild-type cells and n = 802 for 10 LUZP1 knockout cells. Data are represented as mean ± SEM. *, P≤0.05; ***, P≤0.001, Student’s *t* test. (**I**) NM-IIA assemblies in a sucrose density gradient. Identical amounts (100 µg) of total cell lysates from wild type and LUZP1 knockout cells were subjected to sucrose density gradient (10-30%), and total 24 fractions were collected followed by SDS-PAGE and WB with NM-IIA heavy chain antibody. Tubulin was probed as control.

Importantly, LUZP1 deficiency led to a reduction in the phosphorylation levels of Thr18 and Ser19 in the regulatory light chain of NM-II (P-MLC2) (Figure 6B), consistent with a previous finding (Yano et al., 2021). To determine whether the changes in myosin phosphorylation were specifically caused by the loss of LUZP1, a rescue experiment was conducted by introducing exogenous LUZP1-GFP into LUZP1 knockout cells. The results revealed that the deficiencies in NM-II phosphorylation (P-MLC2) could be fully restored through expressing LUZP1-GFP in LUZP1 knockout cells (Figure 6C). Furthermore, the N-terminal extension LUZP1_1-550 was able to rescue the phosphorylation of Thr18/Ser19 in myosin II, similarly to the full-length LUZP1 (Figure 6C). However, the mutant LUZP1_252-550 only partially rescued, and the mutant LUZP1_550-1076 failed to rescue myosin II phosphorylation (Figure 6C). These findings indicate that the N-terminal extension (LUZP1_1-550) is critical for the regulation of myosin II phosphorylation and the assembly of thick contractile actomyosin bundles.

Myosin phosphatase is a critical enzyme involved in myosin phosphorylation, composed of three subunits: a protein phosphatase catalytic subunit PP1c β/δ, a myosin phosphatase targeting subunit MYPT1, and a small regulatory subunit M20 (Ito et al., 2004). Phosphorylation of MYPT1 at Thr696 or Thr853 has been found to inhibit the myosin phosphatase activity, leading to increased phosphorylation of MLC2 (Khromov et al., 2009). To elucidate whether LUZP1 regulates MLC2 phosphorylation via its effect on the phosphorylation of the myosin phosphatase, we examined MYPT1 phosphorylation levels at Thr 853 (P-MYPT1) in both wild type and LUZP1 knockout cells. The results revealed a significant decrease in P-MYPT1 levels in LUZP1 knockout cells (Figures 6D and 6E), suggesting that LUZP1 regulates the phosphorylation of the targeting subunit of the myosin phosphatase MYPT1.

To further understand how LUZP1 regulates the phosphorylation of MLC2, we examined the expression level of ROCK1 in both wild type and LUZP1 knockout cells, as Rho-associated kinase has been identified as the primary player in MLC phosphorylation. The results from Western blot analysis revealed no significant alteration in the ROCK1 abundance upon LUZP1 depletion (Figure 6D). Furthermore, overexpressing ROCK1 in LUZP1-depleted cells failed to rescue the decline in P-MYPT1 caused by LUZP1 depletion, while ROCK1 overexpression restored MLC2 phosphorylation at Ser19 and Thr18 (P-MLC2) to levels similar to those in wild-type cells (Figures 6D and 6E). This suggests that LUZP1 regulates myosin phosphatase phosphorylation via a pathway independent of ROCK1. Together, our data indicate that LUZP1 regulates the phosphorylation of myosin II and the targeting subunit of the myosin phosphatase MYPT1.

### LUZP1 Regulates the Assembly of Large Myosin II Stacks

Given LUZP1’s localization within actin filament bundles containing myosin II and the observed changes in actomyosin organization and contractility upon LUZP1 knockout, it strongly suggests LUZP1’s role in the formation of NM-II structures. To delve into this aspect, we utilized super-resolution microscopy 3D-SIM to assess the structural organization of actomyosin stress fibers in both wild-type and LUZP1 knockout cells. The results revealed that LUZP1 knockout cells exhibited more unipolar NM-IIA assemblies compared to wild-type cells (Figure 6F). Although bipolar NM-IIA filaments could still form in LUZP1 knockout cells, these filaments failed to form larger NM-II stacks, which were abundant in wild-type cells. The average lateral stack length was significantly decreased in LUZP1 knockout cells (Figure 6G). Moreover, the size distribution of NM-IIA stacks changed, with an increased percentage of small NM-IIA stacks in LUZP1 knockout cells (Figure 6H). The reduction of large NMII stacks (diameter >300 nm) in LUZP1 knockout cells indicates that loss of LUZP1 affected the formation of large NM-IIA stacks. This was further supported by results from density-gradient centrifugation (Figure 6I). In wild type cell extracts, NM-IIA was distributed to fractions 2–7, whereas in LUZP1 knockout cell extracts, NM-IIA in the higher sucrose fraction, particularly fraction 3, shifted to fraction 1, corresponding to smaller NM-IIA structures (Figure 6I). Collectively, these findings highlight LUZP1’s critical role in the formation of larger myosin stacks. To validate whether these alterations were specifically attributed to LUZP1 loss, a rescue experiment was conducted by the expression of LUZP1-GFP in LUZP1 knockout cells. The results showed that the defects in the formation of larger NM-II structures in LUZP1 knockout cells can be rescued through LUZP1-GFP expression (Figures 7A and 7B). This underscores LUZP1-GFP’s ability to restore proper stack assembly and myosin II organization in LUZP1 knockout cells. Taken together, these findings demonstrate that LUZP1 deficiency leads to altered actomyosin organization and impaired formation of larger NM-II structures. This suggests that LUZP1 is essential for the formation of larger filament stacks in non-muscle cells.

**Figure 7.**
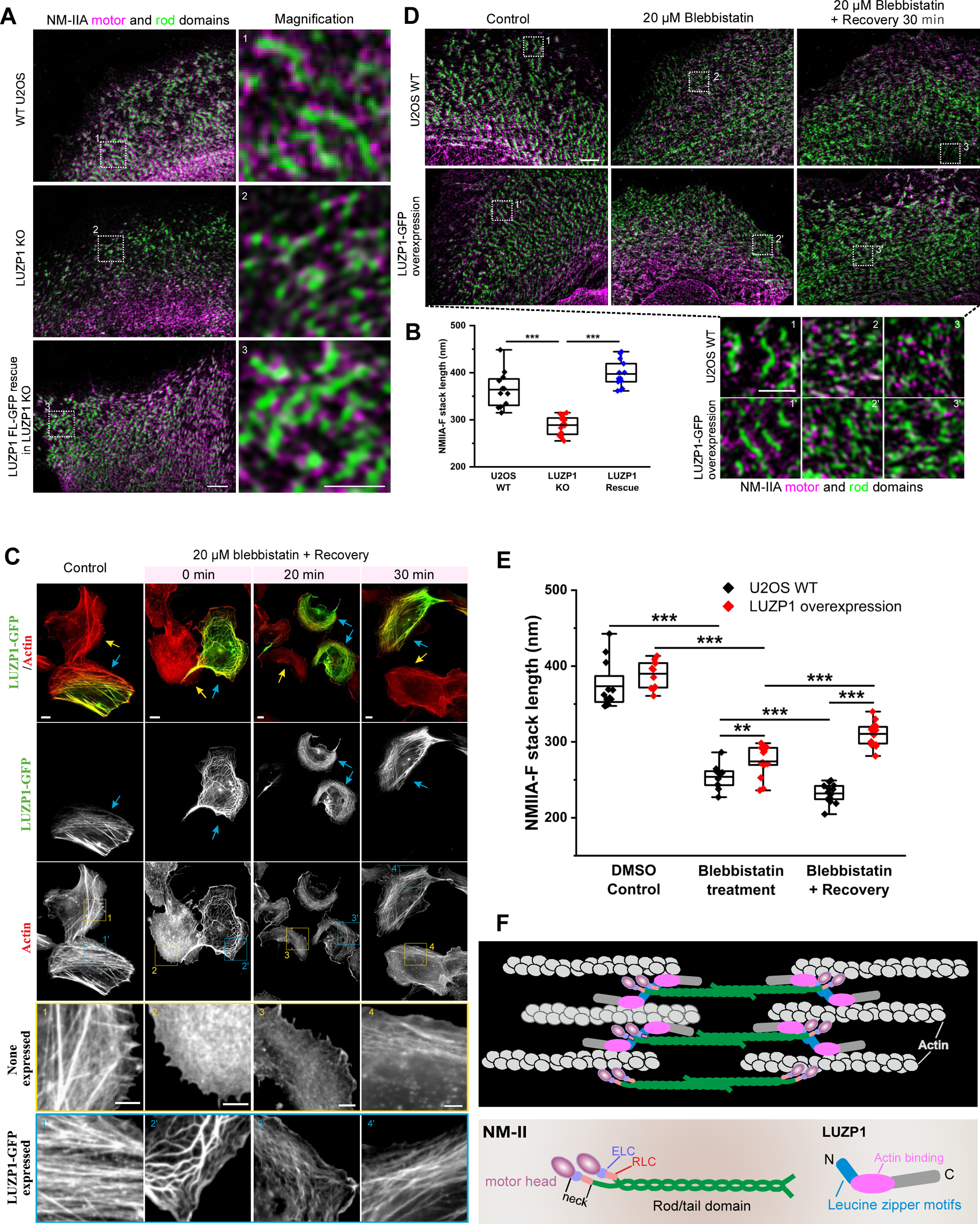
Disruption of NM-II stack assembly LUZP1 knockout cells can rescued by LUZP1 expression in LUZP1 knockout cells. **(A)** Representative 3D-SIM images of myosin stacks in wild type U2OS cells, LUZP1 knockout cells, and LUZP1 knockout cells expressing LUZP1-GFP construct. Cells are stained with NMIIA motor domain antibody (magenta) and NMIIA rod domain antibody (green). The right panels show the magnified regions indicated by boxes with white dotted line. Scale bar: 2 µm, 1 µm for magnified regions. (**B**) Quantification of the average length (µm) of NM-II stacks. n=12 for wild type cells, n = 16 for LUZP1 knockout cells, and n = 13 for LUZP1 knockout cells expressing LUZP1-GFP. Data are represented as mean ± SEM. ***, P≤0.001, Student’s *t* test. **(C)** Representative images of wild-type U2OS cells and wild-type U2OS cells expressing LUZP1-GFP treated with DMSO (DMSO Control), 20 µM blebbistatin for 40 mins (Blebbistatin), and 20 µM blebbistatin for 40 mins followed by recovery in fresh media (Blebbistatin + recovery). Cells were stained with phalloidin (red). Boxes with yellow or blue frames show the magnified regions. Scale bar: 2 µm, 1 µm for magnified regions. (**D**) Quantification of the average length (µm) of NM-II stacks. n = 12 for WT and LUZP1-GFP cells treated with DMSO Control; n = 11 and 15 for WT and LUZP1-GFP cells treated with blebbistatin; n = 13 and 16 for WT and LUZP1-GFP cells treated with blebbistatin followed by recovery. **, P≤0.01, ***, P≤0.001, Student’s *t* test. (**F**) A working model for LUZP1 in the regulation of actomyosin bundle formation. LUZP1 predominantly localizes in close proximity to the neck region of myosin II, where LUZP1 activates NM-II by regulation of NM-II phosphorylation and increases assembly of NM-II stacks. Additionally, LUZP1 interacts with actin and self-oligomerizes to promote the formation of myosin II stacks and actomyosin bundles.

To further examine the correlation between myosin II and LUZP1, we expressed LUZP1-GFP in wild type U2OS cells and treated the cells with blebbistatin, a specific inhibitor of myosin II (Kovács et al., 2004; Straight et al., 2003). After 40 minutes of blebbistatin treatment, transverse arcs, dorsal stress fibers, and most ventral stress fibers were lost in wild-type cells (Figure 7C). However, in cells overexpressing LUZP1, the thick actomyosin bundles were preserved, albeit with a distorted morphology of the actin stress fibers (Figure 7C). This preservation of thick actomyosin bundles in LUZP1-overexpressing cells suggests that LUZP1 might have a protective role in maintaining the structure and organization of stress fibers in the presence of the myosin II inhibitor. This protective effect was further supported by the quicker recovery of actin stress fibers in LUZP1-overexpressing cells after blebbistatin washout. While cells without LUZP1-GFP expression exhibited slow recovery, actin stress fibers returned to their normal morphology in cells expressing LUZP1-GFP after a 30-minute recovery period following blebbistatin treatment (Figure 7C). Consistent with the changes in stress fibers, the average length of NM-IIA stacks significantly decreased after 40 minutes of blebbistatin treatment in both wild-type cells and LUZP1-overexpressing cells compared to cells treated only with DMSO (Figures 7D and 7E). Intriguingly, LUZP1 overexpression showed a notable protective effect in preventing the disruption of NM-IIA stack assembly compared to wild-type cells (Figures 7D and 7E). This protective effect of LUZP1 overexpression was particularly evident after 30 minutes of recovery, where the average length of NM-IIA stacks significantly increased in LUZP1-overexpressing cells following blebbistatin treatment, in contrast to wild-type cells (Figure 7E). These results suggest that LUZP1 has the ability to maintain the integrity of stress fibers against the detrimental effects of the myosin II inhibitor blebbistatin by enhancing NM-IIA stack assembly. In conclusion, these data demonstrate the critical role of LUZP1 in the assembly of large myosin II stacks.

## Discussion

Actin filaments, together with myosin II, assemble into contractile actomyosin bundles that are crucial for adhesion, morphogenesis, and mechanosensing in non-muscle cells. The mechanisms underlying NM-II folding and self-assembly into bipolar filaments as well as their recruitment to the nascent stress fibers have started to emerge (Jiu et al., 2019; Lehtimäki et al., 2017; Sandquist and Means, 2008; Vicente-Manzanares et al., 2007). LUZP is a leucine zipper motif-containing protein that is predominantly expressed in the mouse brain and neural lineages (Gad et al., 2012). The functions of LUZP1 in various cellular contexts are complex and may involve interactions with different components of the cytoskeleton, including microtubules and actin filaments (Bozal-Basterra et al., 2020; Gonçalves, 2022; Gonçalves et al., 2020; Yano et al., 2021). LUZP1 regulates ciliogenesis by interacting with the truncated transcription factor SALL1 (Bozal-Basterra et al., 2020) and the actin-stabilizing protein EPLIN (Gonçalves et al., 2020). Additionally, LUZP1 associates with microtubules at the tight junctions of epithelia cells to promote apical constriction (Yano et al., 2021). The interactions of LUZP1 with actin and actin-associated proteins strongly indicate a crucial role for LUZP1 in the regulation of the actin cytoskeleton (Bozal-Basterra et al., 2020, 2021; Wang and Nakamura, 2019a, 2019b; Yano et al., 2021). However, the intricate molecular details and mechanisms underlying how LUZP1 precisely contributes to stress fiber assembly, maintenance, and contractility in non-muscle cells are not fully understood.

Here, we reveal that LUZP1 associates with actin stress fibers, particularly those enriched in NM-II-containing thick actomyosin bundles. Loss of LUZP1 causes severe defects in the maturation of contractile actomyosin bundles, and consequently leads to abnormal cell migration and a decreased ability to exert forces on the environment. The disruption of these processes in LUZP1 knockout cells provides mechanistic insights into the defective assembly of thick ventral stress fibers and the associated cellular contractility abnormalities. Based on previous research and our data, we propose a working model for the regulation of actomyosin bundle formation by LUZP1 (Figure 7F). LUZP1 predominantly localizes in close proximity to the neck region of myosin II, where LUZP1 activates NM-II by regulation of NM-II phosphorylation and increases assembly of NM-II stacks. Additionally, LUZP1 interacts with actin and self-oligomerizes to promote the formation of myosin II stacks and actomyosin bundles. The N-terminal extension LUZP_1-550 is essential for this role in particular because LUZP_1-550 is required for thick stress-fiber localization and phosphorylation of NM-II. The C-terminal extension localizes only to thin stress fibers and has only a minor role in promoting the formation of thick actomyosin bundles.

In alignment with the previous study (Yano et al., 2021), our finding affirm that LUZP1 plays a critical role in myosin II phosphorylation. Interestingly, prior research indicates that LUZP1 interacts with the catalytic subunit PP1c β/δ of myosin phosphatase and inhibits its activity in vitro (Yano et al., 2021). Our current study contributes an additional layer of understanding by uncovering a novel mechanism through which LUZP1 modulates the phosphorylation of the myosin phosphatase targeting subunit, MYPT1. This newly identified pathway provides insight into how LUZP1’s actions regulate myosin II phosphorylation. Nonetheless, the intricate relationship between the catalytic subunit PP1c beta/delta and the phosphorylation of the myosin phosphatase targeting subunit MYPT1, under the influence of LUZP1, warrants further investigation for a comprehensive understanding.

Importantly, our study demonstrates that LUZP1 is essential for the formation of myosin large stacks, which is closely linked to the assembly and organization of actin filaments within stress fibers. Actin filaments provide the structural framework for stress fibers, while myosin II filaments interact with actin filaments to generate contractile forces. The similarity between the effects of myosin II knockdown and LUZP1 depletion, such as altered cell morphology, decreased F-actin retrograde flow, traction force, and migration speed, further supports the notion that LUZP1 plays a role in regulating myosin II activity and interacts with actin filaments (Cai et al., 2006; Even-Ram et al., 2007; Medeiros et al., 2006) (Figure?). On the other hand, the phenotypes of LUZP1 KO and KD are partially different from those of NMII knockdown. LUZP1 knockout cells displayed a significantly increased number of thin transverse arcs but a diminished number of thick actomyosin bundles. Loss of LUZP1 also resulted in elongated dorsal stress fibers. In contrast, NMII-depleted cells lost most of their stress fibers, including the majority of the dorsal stress fibers and transverse arcs, containing only a few discrete ventral stress fibers attached to focal adhesions at both ends (Even-Ram et al., 2007; Sandquist et al., 2006; Shutova et al., 2017).

The proper assembly and alignment of myosin II filaments within stress fibers are essential for efficient cell migration. A recent study showed that LUZP1 depletion increased cell migration (Bozal-Basterra et al., 2021). In contradiction to their finding, our data revealed that LUZP1 knockout reduced contractile forces and impaired cell migration. Moreover, we showed that siRNA-mediated knockdown of LUZP1 in several cell lines consistently led to impaired cell migration and invasion. Interestingly, the reported phenotypes of cell migration caused by myosin II deficiency are also controversial. Increased or decreased cell migration, or no effects on cell migration, were reported when myosin II was depleted or inhibited (Betapudi et al., 2006; Doyle et al., 2012; Even-Ram et al., 2007; Kim and Adelstein, 2011; Sandquist et al., 2006; Shih and Yamada, 2010; Shutova et al., 2017). Myosin IIA (NMIIA) and myosin IIB (NMIIB) were often present simultaneously in cells, but at different ratios and localizations. NMIIA and NMIIB cooperate for stress fiber formation rather than independently drive the formation of their preferred stress fiber types (Shutova et al., 2017). Given the complexity of myosin II’s functions and its interactions with other cytoskeletal components, it is not surprising to find varying results in different studies. The controversy in the literature could be attributed to several factors, including cell type, the abundance and relative ratio of different myosin isoforms, and compensatory mechanisms activated by cells after myosin depletion. The discrepancy between the previous study and our findings regarding the effect of LUZP1 depletion on cell motility may result from these factors because LUZP1 is involved in the regulation of myosin II phosphorylation, as our and Yano’s work both showed (Yano et al., 2021). The contribution of LUZP1 loss to the phosphorylation of myosin IIA and IIB may vary due to the distinct abundance and relative ratios of myosin IIA and myosin IIB in cells. This highlights the complexity of cellular processes and the need for further investigation to understand the precise mechanisms by which LUZP1 influences cell motility.

Our data also revealed that different regions of LUZP1 play distinct roles in stress fiber assembly and function. The N-terminal region of LUZP1, specifically LUZP1_1-550, is critical for localizing to thick stress fibers and rescuing the defects in actomyosin bundle assembly caused by LUZP1 depletion. Moreover, this region is important for the phosphorylation of the myosin II light chain. Importantly, the leucine zipper motifs at the N-terminus of LUZP1 contributes to the assembly of thick actomyosin bundles because cells expressing LUZP1 mutants LUZP1_252-550 and LUZP1_550-1076, lacking these leucine zipper motifs, only display thin stress fibers. It is known that leucine zipper motifs are essential structural elements that facilitate protein dimerization and, in some cases, oligomerization (McCain and Grytnes, 2010). Through the interactions facilitated by leucine zipper motifs, LUZP1 can create a scaffold that promotes the alignment and bundling of actomyosin filaments (Figure 7C). Dimerization/oligomerization of LUZP1 and its interaction with actin and phosphorylated myosin will enhance the stability and contractile properties of the actomyosin bundles. On the other hand, the C-terminal region of LUZP1, LUZP1_550-1076, fails to rescue the defects in thick actomyosin bundle assembly, despite its co-localization with thin stress fibers. This region of LUZP1 was shown to mediate interactions with other actin-binding proteins such as EPLIN (Gonçalves et al., 2020; Wang and Nakamura, 2019a), suggesting that it may have additional roles in stress fiber regulation through its interactions with other actin-binding proteins. Overall, these findings indicate that different regions of LUZP1 contribute to its cellular function in stress fiber assembly and organization. The N-terminal region is essential for proper localization and function within thick actomyosin bundles, while the C-terminal region may mediates interactions with other actin-binding proteins and fine-tuning stress fiber dynamics.

In conclusion, our study unequivocally establishes LUZP1 as a pivotal factor responsible for orchestrating the assembly and dynamics of stress fibers. LUZP1’s involvement is indispensable for the proper formation of myosin II stacks and the maturation of actomyosin bundles. These functions collectively play a crucial role in upholding the structural integrity and optimal functionality of the actin cytoskeleton.

## Materials and Methods

### Materials

Polyclonal rabbit antibody against LUZP1 (#HPA028542, 1:50 dilution for IF analysis; 1:500 dilution for WB), monoclonal mouse antibody (#V9131) against full-length human vinculin, (1:400 IF), mouse monoclonal (M4401) against NMIIA-RLC (1:100 IF), mouse monoclonal antibody against α-tubulin C-terminus (#T5168, 1:10,000 WB) and mouse monoclonal antibody recognizing the N-terminus β-Actin (#A5441, 1:10,000 WB) were purchased from Merck. Rabbit polyclonal phosphomyosin (Thr18/Ser19) RLC (#3674, 1:1000 dilution for WB) was purchased from Cell Signaling Technology. Rabbit polyclonal antibody (#PAB1195) recognizing the full-length GAPDH (1:10,000 WB) was purchased from Abcam. Polyclonal NMHC-IIA antibody (#909801) binding to the C-terminal tail residues 1948-1960 (1:1,000 IF and WB) and NMHC-IIB antibody (#909901) binding to residues 1965-1976 in the C terminus (1:100 IF, 1:400 WB) were purchased from BioLegend. Rabbit polyclonal antibody against GFP tag (#50430-2-AP, 1:4,000 WB) was purchased from Proteintech. DAPI (#D1306; 1:10,000 IF). Alexa fluor phalloidin 568 (#A12380, 1:100 IF) and 647 (#A22287, 1:100 IF), Alexa Fluor goat anti-mouse 488 (#A11001), 568 (#A11004), and 647 (#A31571), Alexa Fluor goat anti-rabbit 488 (#A11034, 1:200 IF) and 647 (#A21245, 1:200 IF), and both HRP-conjugated goat anti-rabbit (#G-21234, 1:20, 000) and HRP-conjugated goat anti-mouse (G-21040; 1:10,000) antibodies were purchased from Jackson ImmunoResearch.

### CRISPR construct design

A guide sequence targeting exon 1 of the human *LUZP1* gene was selected based on the CRISPR Design Tool (Ran et al., 2013). Oligonucleotides for cloning guide RNA into the pSpCas9 (BB)-2A-GFP vector (48138; a gift from F. Zhang, Addgene, Cambridge, MA) were designed as described previously (Ran et al., 2013). Transfected cells were detached 24 hours after transfection and suspended in complete DMEM supplemented with 10 mM Hepes. Cells were subsequently sorted with FACSAria II (BD), using low-intensity GFP-positive pass gating. A GFP-positive single cell was sorted into one well of a 96-well plate containing 200 μL of DMEM containing 20% FBS and 10 mM Hepes. CRISPR clones were cultivated for approximately 2 weeks before selecting clones with no discernible LUZP1 protein expression. Five clones exhibiting identical phenotypes with no detectable LUZP1 protein expression survived. One clone was selected for use in this study, and the LUZP1 knockout was validated with sequencing. In order to confirm the CRISPR knockout, several primers were designed around the guide RNA binding site. Only primers that were more than 2 kbp away from the site gave the PCR product, indicating that there is a big deletion. The PCR products were confirmed by Sanger sequencing. The region for sequencing was amplified using primers: forward primer (LUZP1_NGS_F): ACACTCTTTCCCTACACGACGCTCTTCCGATCTTGATGATGGTCTCCAGCT GT; Reverse primer (LUZP1_NGS_R4): AGGCATTCCGCACAGTAACA. The obtained PCR products were sanger-sequenced with LUZP1_NGS_F primer.

### Plasmids

The Full-length human LUZP1 coding sequence in the pENTR221 vector (ORFeome Library, Genome Biology Unit supported by HiLIFE and the Faculty of Medicine, University of Helsinki, and Biocenter Finland) was cloned into the pEGFP-N1 backbone (Takara Bio Inc.) with XhoI and Kpn1 restriction sites. Two missense mutations were corrected by cloning. The truncated mutants were cloned from full-length LUZP1-GFP using the same restriction sites. mEmerald-MyosinIIA-N-14 (Addgene plasmid #54191) and mApple-LC-myosin-C-10 (Addgene plasmid #54919) were gifts from Michael Davidson. The human GFP-β-actin plasmid was a gift from M. Bahler (Westfalian Wilhelms-University, Münster, Germany). Full-length human ROCK1 coding sequence in pDONR221 (ORFeome Library, Genome Biology Unit supported by HiLIFE and the Faculty of Medicine, University of Helsinki, and Biocenter Finland) was cloned into a destination vector tagged with EGFP.

### Cell Culture

Human osteosarcoma (U2OS) cells and human ovarian cancer cell line A2780 were maintained in high-glucose (4.5 g/L) DMEM, supplemented with 10% FBS (Gibco), 10 U/ml penicillin, 10 µg/ml streptomycin, and 20 mM **L**-glutamine (10378-016; Gibco). Human breast cancer cell line MDA-MB-231 was maintained in RPMI-1640, supplemented with 10% FBS (Gibco), 10 U/ml penicillin, 10 µg/ml streptomycin, and 20 mM **L**-glutamine (10378-016; Gibco). All cells were regularly tested negative for mycoplasma with MycoAlert Plus (Lonza) and maintained at 37°C in a humidified atmosphere with 5% CO_2_ flow. Transient transfections were performed with Fugene HD (#E2311, Promega), according to the manufacturer’s instructions using a 3.5:1 Fugene/DNA ratio and 48 hours of incubation before fixation with 4% PFA in PBS. For live cell and SIM imaging, cells were detached 24 h after transfection with Trypsin-EDTA (0.25% wt/vol) and re-plated onto 10 µg/ml fibronectin (Roche) pre-coated MatTek dishes with glass bottoms. siRNA was transfected with lipofectamine RNAiMAX (#13778-075, Thermo Fisher), according to the manufacturer’s instructions, using 30 nM Hs_LUZP1 siRNA (target sequence, 5′-CAGCGGGTGCTGAGAATTGAA-3′; QIAGEN) or 30 nM AllStars negative-control siRNA (QIAGEN). A 72-hour incubation period was used to deplete the target protein.

### Western blotting

U2OS cells were lysed for 10 min at 4°C with radioimmunoprecipitation assay (RIPA) lysis buffer (50 mM Tris (pH 7.4), 150 mM NaCl, 1% NP-40, 0.5% sodium deoxycholate, 1 mM PMSF, 10 mM DTT, 40 μg/ml DNaseI mM, 1 mM Na3VO4 and 1 µg/ml aprotinin, leupeptin). The lysates were briefly sonicated before centrifugation at 13,000 rpm for 10 min at 4°C. Protein concentrations were determined with Bradford reagent (500-0006; Bio-Rad Laboratories) and equal amounts of protein were loaded and run on 4-20% gradient SDS-PAGE gels (4561096; Bio-Rad Laboratories). Proteins were transferred from SDS-PAGE gels to nitrocellulose membranes using Trans-Blot Turbo (Bio-Rad Laboratories). Membrane was blocked in 5% milk or 5% BSA in TBS and incubated overnight at 4°C and for 1 hour at RT with primary and secondary HRP-conjugated antibodies, respectively. Proteins were detected with Western Lightning ECL Pro substrate (PerkinElmer).

### Real-time quantitative PCR

Total mRNA was extracted with the GeneJET RNA purification kit (K0731; Thermo Fisher Scientific) and single-stranded cDNA was synthesized (K1671; Thermo Fisher Scientific) from 500 ng of the extracted mRNA. Primers amplifying around 200-bp region from the isoforms of human LUZP1 (LUZP1-1076aa and LUZP1-1026aa), and human GAPDH were designed with the Primer-BLAST tool (Ye et al., 2012). Real-time quantitative PCR (RT-qPCR) reactions were performed with Maxima SYBR Green/ROX (K0221; Thermo Fisher Scientific) in Bio-Rad Laboratories CFX96. mRNA levels were calculated with the 2^−ΔΔCt^ method, normalized to GAPDH (ΔCt) and WT expression levels, respectively.

### Primers used in this study

Both forward (F) and reverse (R) primers used in this study are presented. Complementary sequences to the target cDNAs are underlined.

Guide RNA oligonucleotides for CRISPR, targeting exon 1 of human LUZP1. **e1-hsLUZP1-F**: 5*′*-CACCGGTAGCTTAAACCGCAAGTGG-3*′*, **e1-hsLUZP1-R**: 5*′*-AAACCCACTTGCGGTTTAA GCTACC-3*′*.

Cloning primers for LUZP1-1076aa-GFP (LUZP1-GFP) and truncated GFP tagged LUZP1-1076aa proteins: **LUZP1-GFP-F**: 5*′*-CGCGCTCGAGATGGCCGAATTTACAAGCTACAAG-3*′*, **LUZP1-GFP-R**: 5*′*-CGCGGGTACCACGTTCTCCTCAGCACAGGGCCTGGT-3*′*; **LUZP1_1-252-F**: 5*′*-CA CTGCCGTCCACGGTACCGCGGGCCCG-3*′*, **LUZP1_1-252-R**: 5*′*-GGTACCGTGGACG GCAGTGTGGAAGAGATGCC-3*′*; **LUZP1_1-550-F**: 5*′*-GGCAACGGAAGTACGGTACCGCGGGCCCG-3*′*, **LUZP1_1-550-R**: 5*′*-GGTACCGTACTTCCGTTGCCAAGCACGTG TC-3*′*; **LUZP1_550-1076-F**: 5*′*-GATCTCGAGATGAGTCAAGTAACTCAGGCTGCAAA C-3*′*, **LUZP1_ 550-1076-R**: 5*′*-TACTTGACTCATCTCGAGATCTGAGTCCGGTAGC-3*′*; **LUZP1_252-550-F**: 5*′*-CGCGCTCGAGATGAAAGAATCAAGAAGGAAGGGTGG-3*′*, **LUZP1_252-550-R**: 5*′*-CGCGGGTACCGTACTTCCGTTGCCAAGCACGTG-3*′*.

Quantitative PCR oligonucleotides for LUZP1_1076aa, LUZP1_1026aa and human GAPDH. **LUZP1 _1076aa-F**: 5*′*-CTCAGCAAGCATGGAGGAAG-3*′*, **LUZP1_1076aa-R**: 5*′*-AGGCAGTTCAGACG GATCCA-3*′*; **LUZP1_1026aa-F**: 5*′*-CTCACTGTGTCAGAGGTGCT-3*′*, **LUZP1_1026aa-R**: 5*′*-GC TGCTCATGCTTGCTGAGT-3*′*; **GAPDH-F**: 5′-TGCACCACCAACTGCTTAGC-3′, **GAPDH-R**: 5′-GGCATGGACTGTGGTCATGAG-3′.

### Immunofluorescence microscopy

U2OS cells were fixed with 4% PFA for 20 min at RT, followed by three washings with PBS, and permeabilized with 0.1% Triton X-100 in PBS for 8 min. The permeabilized cells were subsequently blocked with 0.2% BSA in Dulbecco’s PBS. Primary and secondary antibodies diluted with Dulbecco’s PBS were incubated with cells overnight at 4°C and 1 h at RT in a dark, humidified chamber, respectively. Alexa-conjugated phalloidin was added along with primary antibodies. For normal immunofluorescence (IF) imaging, samples were directly mounted in Mowiol, containing 2.5% wt/vol 1,4-diazabicyclo [2.2.2] octane (DABCO, D27802, Sigma-Aldrich). Samples for 3D-structured illumination microscopy (3D-SIM) were prepared according to a previous study (Kraus et al., 2017). Briefly, cells were further post-fixed with 4% PFA for 10 min at RT after the secondary antibody incubation. After washing three times with PBS, samples were mounted in Prolong™ Glass Antifade Mountant (P36980, Thermo Fisher Scientific). All IF data were obtained with a Leica TCS SP8 laser scanning confocal microscope with a 63 x 1.3 NA glycerol objective. The numbers and widths of actin filament bundles from cells plated on crossbow-shaped micro-patterns were analyzed using FilamentSensor software as described before (Eltzner et al., 2015). The lamella width, cell height, and nuclei position on crossbow-shaped micro-patterns were analyzed by Image J (version 1.53c).

### 3D-SIM

All 3D-SIM images were obtained at RT using Deltavision OMX Super Resolution (Cytiva) with a 60 x Plan-Apochromat N/1.42 NA oil objective, 1.516 RI immersion oil, a laser module with 488-, 568-, and 640 nm diode laser lines, and three sCMOS cameras, and operated through Acquire SR 4.4 acquisition software. SI reconstruction and image alignment were performed with SoftWoRx 7.0. Imaging arrays of 1024 x 1024 or 512 x 512 were used, both with pixel sizes of 0.08 μm, 0.08 μm, and 0.125 μm (x, y and z).

### Live cell imaging

For measuring the flow rate of arc filaments, U2OS cells were trypsinized after being transiently transfected with GFP-actin for 24 h and re-plated prior to imaging on 10 μg/ml fibronectin-coated glass-bottomed dishes (MatTek Corporation). The time-lapse live cell images were acquired with a Zeiss LSM 880 confocal microscope combined with an Airyscan detector. ZEN software (Zeiss), a 63 x magnified plan-apochromat oil immersive objective with NA=1.40 were used for the image acquisition. Culture dishes were placed in a 37°C sample chamber with a controlled 5% CO2 flow. The recording setting is every 5 seconds for at least 15 minutes. One focal plane was recorded for all the time-lapse videos. The flow rate of arc filaments was measured by manual blind quantification from the same frame of the time-lapse videos.

### Migration and Invasion Assay

For random cell migration measurement, cells were plated on 10 μg/ml fibronectin-coated twelve-well plates (Greiner), and the plate lid was switched to Cell-Secure (CM Technologies) enabling the insertion of CO_2_ input and output valves. Cells were allowed to attach for two hours, washed once with PBS, and replaced with complete DMEM containing 10 mM HEPES prior to starting imaging. Phase contrast time-lapse imaging of migrating cells was conducted on the continuous cell culturing platform Cell-IQ (CM Technologies). The average migration velocity of wild-type and LUZP1 knockout cells was quantified by tracking the nucleus movement in between 20 min imaging cycles for 12 hours with a Cell-IQ analyzer (CM Technologies). Only cells that did not collide with one another were selected for measurements. Mean squared displacement (MSD) is calculated according to the following formulas:

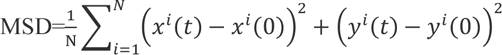

For the scratch wound-healing assay, U2OS and LUZP1 knockout cell monolayers with 90-95% confluent were scratched with a 200-μL pipette tip, resulting in a cell-free gap between two adjoining areas. Cells were washed three times with PBS to remove cell debris, and complete DMEM supplemented with 10 mM HEPES was added. Cells that migrated into the wound area were recorded with every 2 h imaging cycle for 12 hours using the continuous cell culturing platform Cell-IQ (CM Technologies). The gap distance was measured using image J (version 1.53c).

For cell migration and invasion, 1–2×10^5^ cells in 200 μl of serum-free medium were seeded into individual wells of an 8.0 mm, 24-well plate chamber insert (354578, Corning Life Sciences). Then, 500 ul of culture medium containing 10% FBS was added to the bottom of the insert, and the plate was incubated at 37 ^◦^C with 5% CO_2_ for 24 h. After removing the medium, the inserts were washed with PBS, and the cells on the upper surface of the insert were gently removed by scrubbing with a cotton swab. The cells on the bottom surface of the insert were then sequentially fixed with 4% PFA for 5 min, stained with 0.5% crystal violet blue for 5 min, washed three times with PBS, and rinsed with double-distilled water. The positively stained cells were observed using a fluorescence microscope. For the cell invasion assay, Matrigel-coated chambers (354483, Corning Life Sciences) were utilized instead of the chamber inserts used in the migration assay.

### Traction force microscopy

To measure the actomyosin forces that cells exert on their underlying substrate, we used traction force imaging as previously described (Jiu et al., 2017). In brief, both wild-type and LUZP1 knockout cells were cultured for 3-8 hours on custom made 35 mm dishes (Matrigen) coated with fibronectin and displaying specific stiffness (Young’s modulus = 25 kPa). Cells and the underlying microspheres were imaged with a 3I Marianas imaging system containing a heated sample chamber (+37°C) with 5% CO_2_ flow (3I intelligent Imaging Innovations, Germany). 63x/1.2 W C-Apochromat Corr WD = 0.28 M27 objective was used. After the first set of images, cells were detached from the substrates with 2.5% Trypsin (Lonza) and a second set of microsphere images were taken to serve as reference images. Displacement maps were achieved by comparing the reference microsphere images to the first set of images, and by knowing the displacement field and substrate stiffness, cell-exerted traction fields were computed by Fourier-Transform Traction Cytometry. Root mean squared (RMS) magnitudes were computed from the traction fields. Analyses were performed blind, and cell borders were manually traced.

### Density-gradient fractionation

NM-IIA fractionation was performed as previously described with slight modifications (Shutova et al., 2012). In brief, cells were scraped from 125-mm cell culture dishes into lysis buffer, and the supernatant was collected after centrifuging for 20 min at 20,000 g at 4 °C. The amounts of proteins were measured with a Bradford assay, and equal amounts (900 µg) of proteins were applied on the top of a 16 ml 10-30% continuous sucrose density gradient and centrifuged at 74,400 g for 15 h at 4 °C (SW 32.1 Ti; Beckman Coulter). 24 equal volume fractions were collected, precipitated, and resuspended into the Laemmli sample buffer. The samples were separated with SDS-PAGE, transferred to nitrocellulose membranes, and probed with NMHC-IIA and α-tubulin antibodies.

### Statistical Analysis

Statistics were performed with Excel (Microsoft). Sample sizes and the number of replications are indicated in the figures. For data following a normal distribution, Student’s two-sample unpaired t test was used. If the data did not follow a normal distribution, Mann-Whitney u-test for two independent samples was conducted. For all normalizations to WT expression/protein levels, the mean value obtained from the WT sample was set to 1, and the individual intensity data was normalized to that value. For quantitative PCR, the relative gene expression was calculated by the 2^−ΔΔCt^ method. Both WT and knockout/knockdown data were normalized to GAPDH before further normalization. Statistical differences in RMS traction between the WT and LUZP1 KO groups were assessed by using the non-parametric Mann-Whitney-Wilcoxon rank-sum (MWW) test. Geneious (Biomatters Limited) analysis tool was used for sequence alignments in supplemental Figure S3C.

The quantification of NM-IIA filament population distribution from the SIM data was conducted as described in (Fenix et al., 2016) with the following modifications: 5-µm-wide area of the lamellum was drawn with ImageJ (version 1.53c), starting from the most distal NM-IIA filament detected along the cell edge. The NM-IIA stack length in the stress fibers was manually measured using ImageJ.

For the quantification of focal adhesion, all images were manually and blindly quantified from control and LUZP1 depleted cells with ImageJ. Each individual adhesion size was measured with the ROI Manager and freehand line tools in ImageJ. Cells that adhered to several neighbors were discarded from the analysis. Adhesion was classified into seven size groups, and the percentual ratio of the focal adhesion in each class was obtained by dividing the focal adhesion number of individual size classes with the total number of focal adhesions.

## Acknowledgements

Special thanks go to Prof. Pekka Lappalainen for his kindness in providing some of the reagents used in this study and for his critical comments and valuable suggestions for the manuscript. Tommi Kotila is acknowledged for his technical support. We thank Dr. Benjamin Eltzner for his assistance with the FilamentSensor analysis. We thank the DNA Dream Lab facility at the HiLIFE, Institute of Biotechnology for its design and cloning to correct the two missense mutations in LUZP1. We thank the Biocentrum Helsinki Genome Biology Unit for providing gene constructs. We are grateful to the Light and Electron Microscopy Unit (Biocenter Finland), HiLIFE Light Microscopy Unit in Viikki, and Biomedicum Imaging Unit at the University of Helsinki for their kind service in imaging. This study was supported by grants from the Academy of Finland (H.Z. decision No.323670), Jane and Aatos Erkko Foundation (H.Z.), and Guangxi distinguished expert funding (H.Z.).

## Author contributions

H.Z. conceived the project and performed the expression of wild-type and truncated LUZP1 proteins in wild-type and LUZP1 knockout cells. Z.Y. cloned most constructs, purified recombinant proteins, and performed the GTPase assay. S.T. performed the traction force data analysis together with L.W. and H.Z. H.Y.T. performed imaging by superresolution microscopy, analysis of the localization of LUZP1, analysis of myosin stack length, and ROCK1 rescue experiment. C.K. imaged the co-localization of LUZP1 with the myosin light chain by superresolution microcopy. K.K. cloned the LUZP1 constructs for actin binding assays and designed sequencing analysis of the LUZP1 knockout cells together with L.W. X.L. cloned the ROCK1 construct. L.W. carried out all the rest of the experiments and data analysis. H.Z. wrote the manuscript with input from L.W. and H.Y.T. The authors declare no competing financial interests.

## Notes

### Competing Interest Statement

The authors have declared no competing interest.

### Summary of Updates

The abstract and contents of the results have been revised.

