## Supplementary material for "LUZP1 regulates the assembly of stress fibers by promoting maturation of contractile actomyosin bundles": Supplemetary Figures

#### Supplementary Figures

**Figure S1. The cellular localization of GFP-tagged LUZP1 in U2OS cells.** (**A**) Representative images of U2OS cells expressing different LUZP1 isoforms, 1076aa-GFP and 1026aa-GFP. The actin filaments were visualized by Alexa Fluor 568 phalloidin. Scale bar: 10 µm. (**B**) Comparison of cellular localization of endogenous LUZP1 (antibody staining) with LUZP1-1076aa-GFP in U2OS cells. Scale bar: 10 µm. The yellow arrows indicate the bright puncta.

**Figure S2. Contribution of different LUZP1 domains to its cellular localization. (A)** Schematic diagram of domain structures of LUZP1. **(B)** Representative images of cells expressing different GFP-tagged LUZP1 domains. NMIIA was stained with antibody (Alexa Fluor 568) and actin was stained with Alexa Fluor 647 phalloidin.

**Figure S3. Co-localization analysis of LUZP1 with α-actinin and the regulatory light chain of myosin II.** (**A**) Co-localization analysis of LUZP1 with α-actinin. α-actinin (red) and LUZP1 (green) were visualized by antibodies against α-actinin and LUZP1, respectively. Scale bar: 10 µm. Magnified images (corresponding to the orange box) shows the cellular distributions of α-actinin and LUZP1. High-magnification images of individual filaments (corresponding to the white boxes in the magnified regions) illustrate that LUZP1 does not co-localizes with α-actinin. The α-actinin and LUZP1 were indicated with red and green arrows, respectively. The line profiles (corresponding to the white line in the magnified regions) illustrate the distribution of LUZP1 and α-actinin. (**B**) Co-localization analysis of LUZP1 with the regulatory light chain of myosin II. The regulatory light chain (RLC) of myosin II (red) and LUZP1 (green) were visualized by antibodies against the RLC and LUZP1, respectively. Scale bars, 10 and 2 µm in the left and right panels, respectively. In the schematic diagram of NM-II, ELC indicates the essential light chain. The line profiles (corresponding to the white line in the magnified regions) illustrate the distribution of LUZP1 and RLC.

**Figure S4. Validation of LUZP1 knockdown and knockout in U2OS cells.** (**A**) Real-time quantitative PCR (RT-qPCR) analysis of the relative silencing efficiency of LUZP1 isoforms at the transcription level, after treatment with LUZP1 specific siRNA for 72 h. (**B**) Western blot analysis of endogenous LUZP1 protein levels after 72h treatment with control or LUZP1 target-specific siRNA in U2OS cells, as well as the LUZP1 protein levels in wild-type U2OS and LUZP1 knockout cells. GAPDH was probed for equal sample loading. (**C**) The agarose gel electrophoresis of PCR-amplified products from wild-type and LUZP1 knockout U2OS cells. (**D**) Sequencing analysis of

PCR-amplified products in the (C) panel. The gRNA, indicated by a magenta arrow, caused a deletion of 2079 bp nucleotides in the *LUZP1* gene in the knockout cells.

**Figure S5. LUZP1 knockout and knockdown affected the maturation of stress fibers.** (**A & B**) Representative images of wild-type and LUZP1 knockout/knockdown cells grown on fibronectin coated cover glass (A) and crossbow shaped micropatterns (B). The actin filaments were visualized by Alexa Fluor 568 phalloidin and the endogenous LUZP1 was stained using the LUZP1 antibody. Scale bar: 10 µm. (**C**) Representative images of control and LUZP1 knockout cells presenting the effects of LUZP1 knockout on actin filaments, NM-IIA, and vinculin visualized by fluorescence phalloidin and antibodies. The wild-type and LUZP1 knockout cells were grown on fibronectin coated cover glasses with micropattern. Scale bars, 10 µm.

**Figure S6. Effects of LUZP1 knockout on the abundance of NM-IIB in stress fibers.** (**A**) Representative images of actin and myosin II visualized by fluorescent phalloidin and NM-IIB heavy chain antibody, respectively. The wild-type and LUZP1 knockout cells were grown on fibronectin coated cover glasses. Scale bars, 10 µm. The myosin intensity was normalized to the actin intensity (right panel). n = 105 for wild-type cells and n = 123 for LUZP1 knockout cells.

**Video S1. Time-lapse video of transverse arc centripetal flow visualized in wild-type U2OS cells expressing GFP-Actin**. The time-lapse movie was obtained for 200 frames with 5 s intervals using Zeiss LSM 880 confocal microscope. The display rate is 7 frames per second.

**Video S2. Time-lapse video of transverse arc centripetal flow visualized in LUZP1 knockout U2OS cells expressing GFP-Actin**. The time-lapse movie was obtained for 200 frames with 5 s intervals using Zeiss LSM 880 confocal microscope. The display rate is 7 frames per second.

### Figure S1

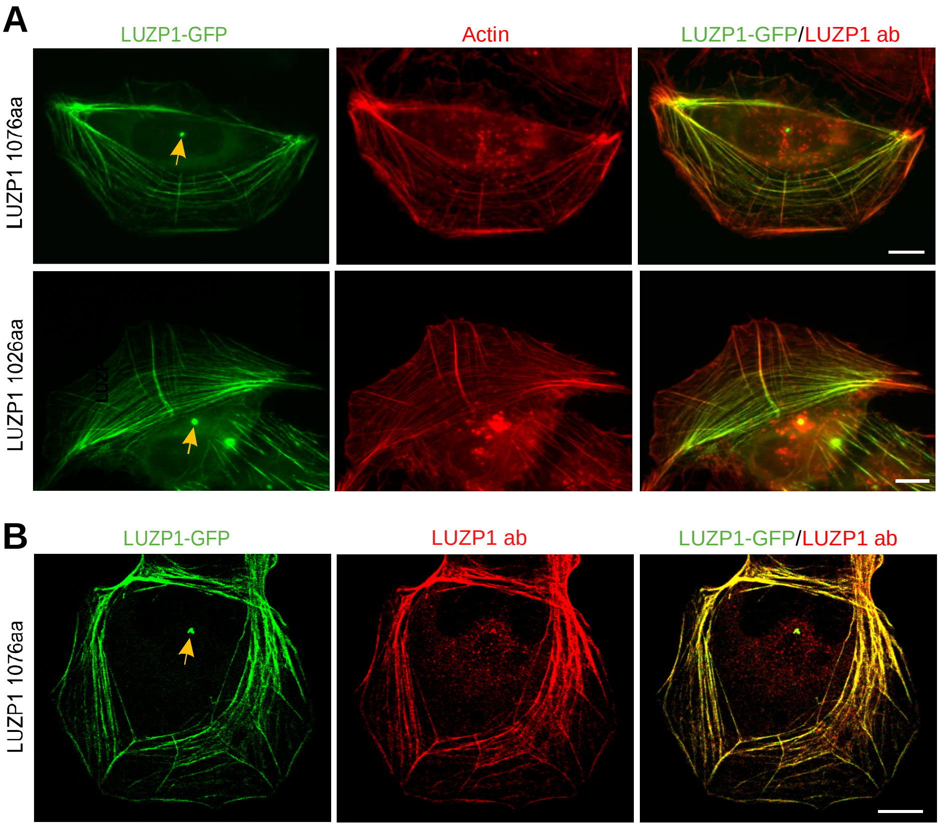
A
LUZP1-GFP
Actin
LUZP1-GFP/LUZP1 ab
LUZP1 1076aa
LUZP1 1026aa
B
LUZP1-GFP
LUZP1 ab
LUZP1-GFP/LUZP1 ab
LUZP1 1076aa

### Figure S2

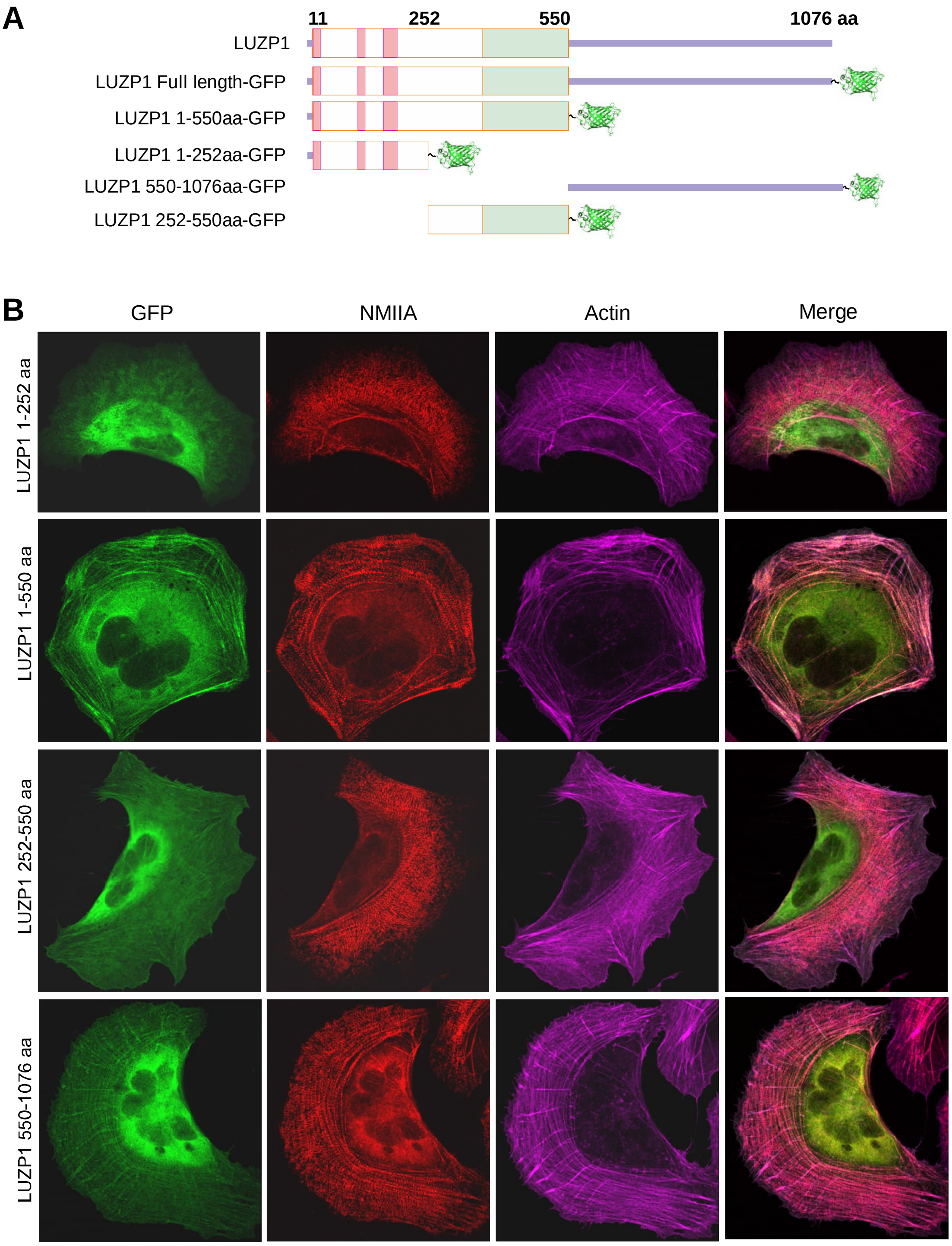

A
11
252
550
1076 aa
LUZP1
LUZP1 Full length-GFP
LUZP1 1-550aa-GFP
LUZP1 1-252aa-GFP
LUZP1 550-1076aa-GFP
LUZP1 252-550aa-GFP
B
GFP
NMIIA
Actin
Merge
LUZP1 1-252 aa
LUZP1 1-550 aa
LUZP1 252-550 aa
LUZP1 550-1076 aa

### Figure S3

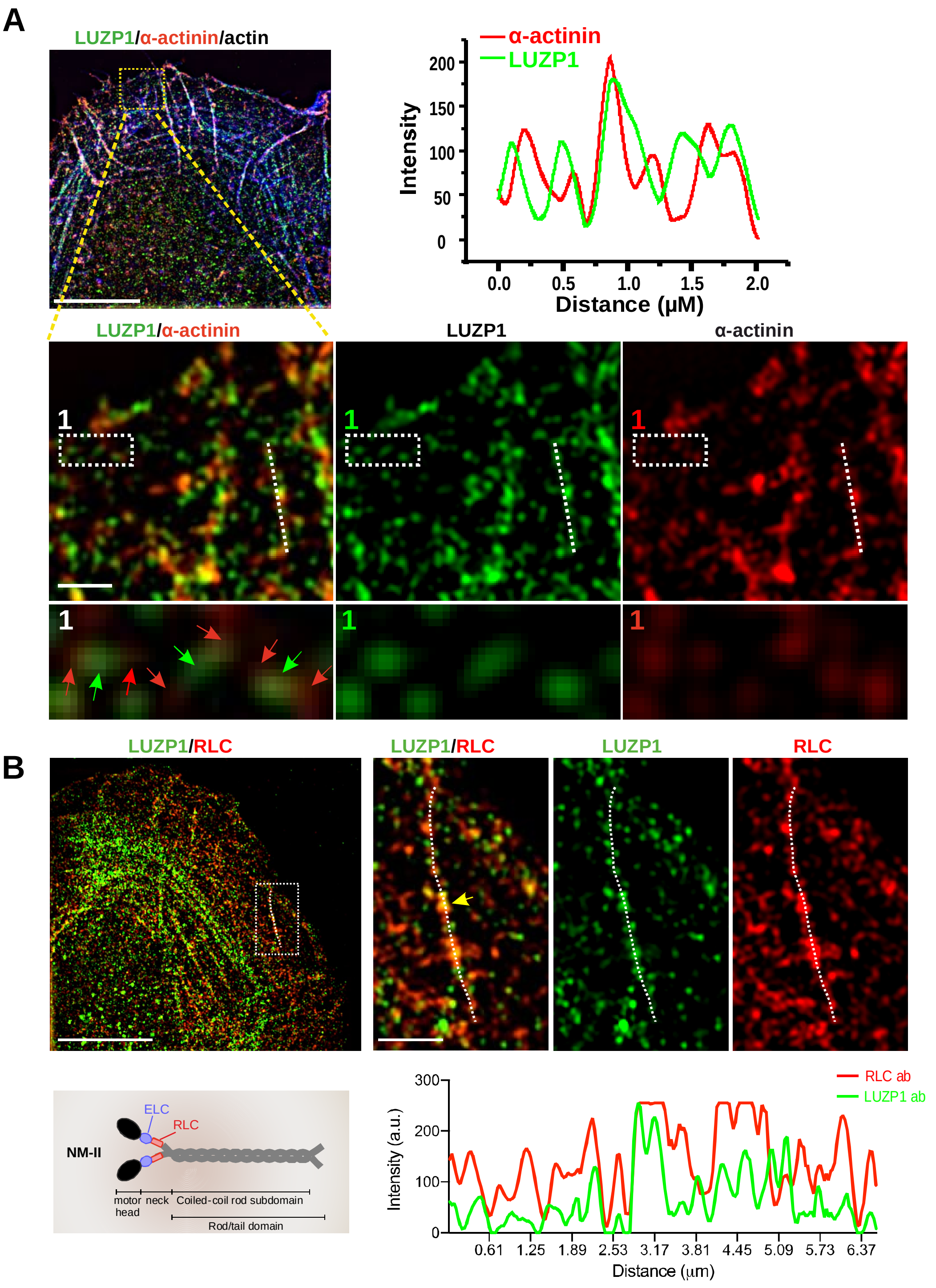
A
LUZP1/α-actinin/actin
α-actinin
LUZP1
Intensity
200
150
100
50
0
0.0
0.5
1.0
1.5
2.0
Distance (μM)
LUZP1/α-actinin
LUZP1
α-actinin
1
B
LUZP1/RLC
LUZP1/RLC
LUZP1
RLC
ELC
RLC
NM-II
motor head
neck
Coiled-coil rod subdomain
Rod/tail domain
300
200
100
0
Intensity (a.u.)
0.61
1.25
1.89
2.53
3.17
3.81
4.45
5.09
5.73
6.37
Distance (μm)
RLC ab
LUZP1 ab

### Figure S4

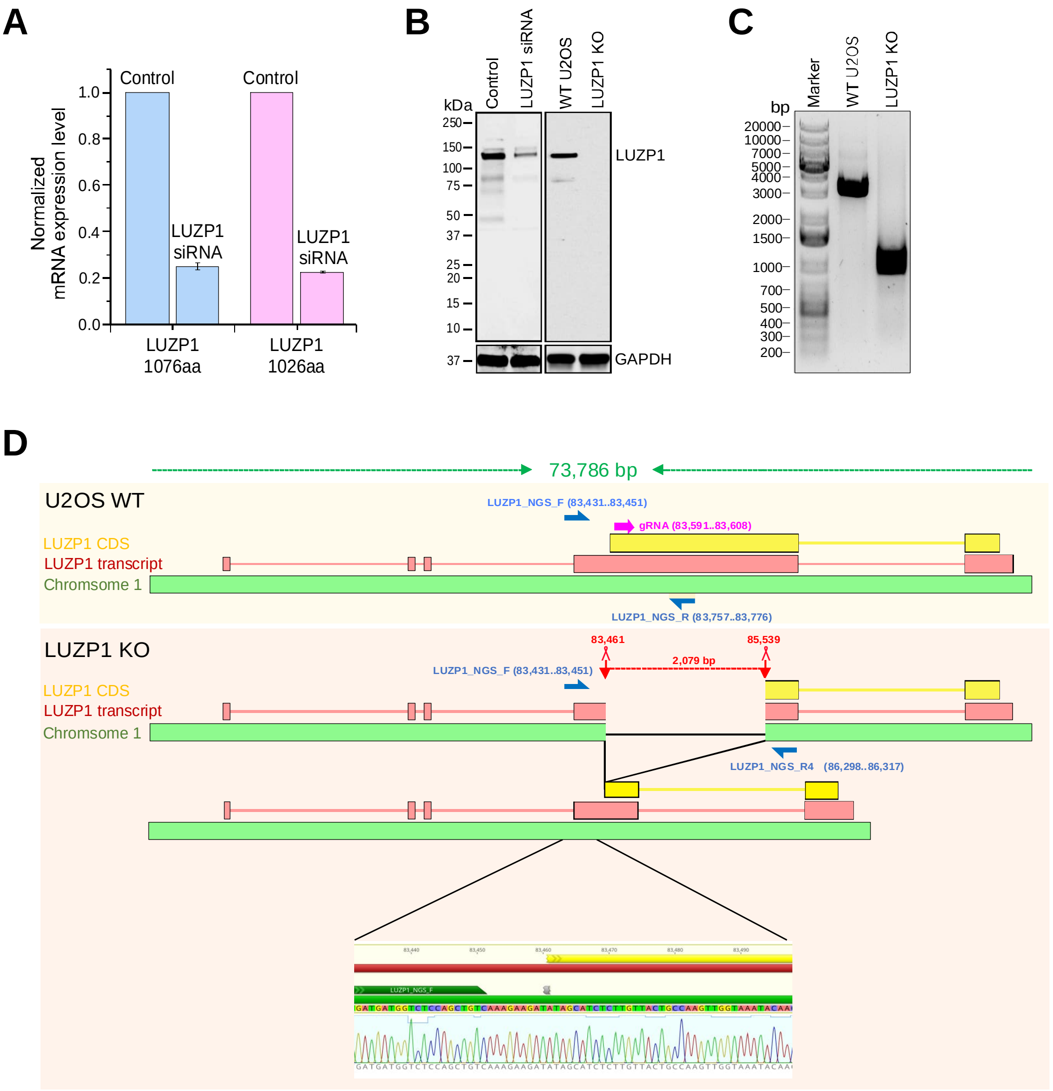
A
Normalized mRNA expression level
Control
LUZP1 siRNA
Control
LUZP1 siRNA
LUZP1 1076aa
LUZP1 1026aa
B
kDa
Control
LUZP1 siRNA
WT U2OS
LUZP1 KO
LUZP1
GAPDH
C
bp
Marker
WT U2OS
LUZP1 KO
D
73,786 bp
U2OS WT
LUZP1_NGS_F (83,431..83,451)
gRNA (83,591..83,608)
LUZP1 CDS
LUZP1 transcript
Chromsome 1
LUZP1_NGS_R (83,757..83,776)
LUZP1 KO
83,461
85,539
2,079 bp
LUZP1_NGS_F (83,431..83,451)
LUZP1 CDS
LUZP1 transcript
Chromsome 1
LUZP1_NGS_R4 (86,298..86,317)

-

### Figure S5

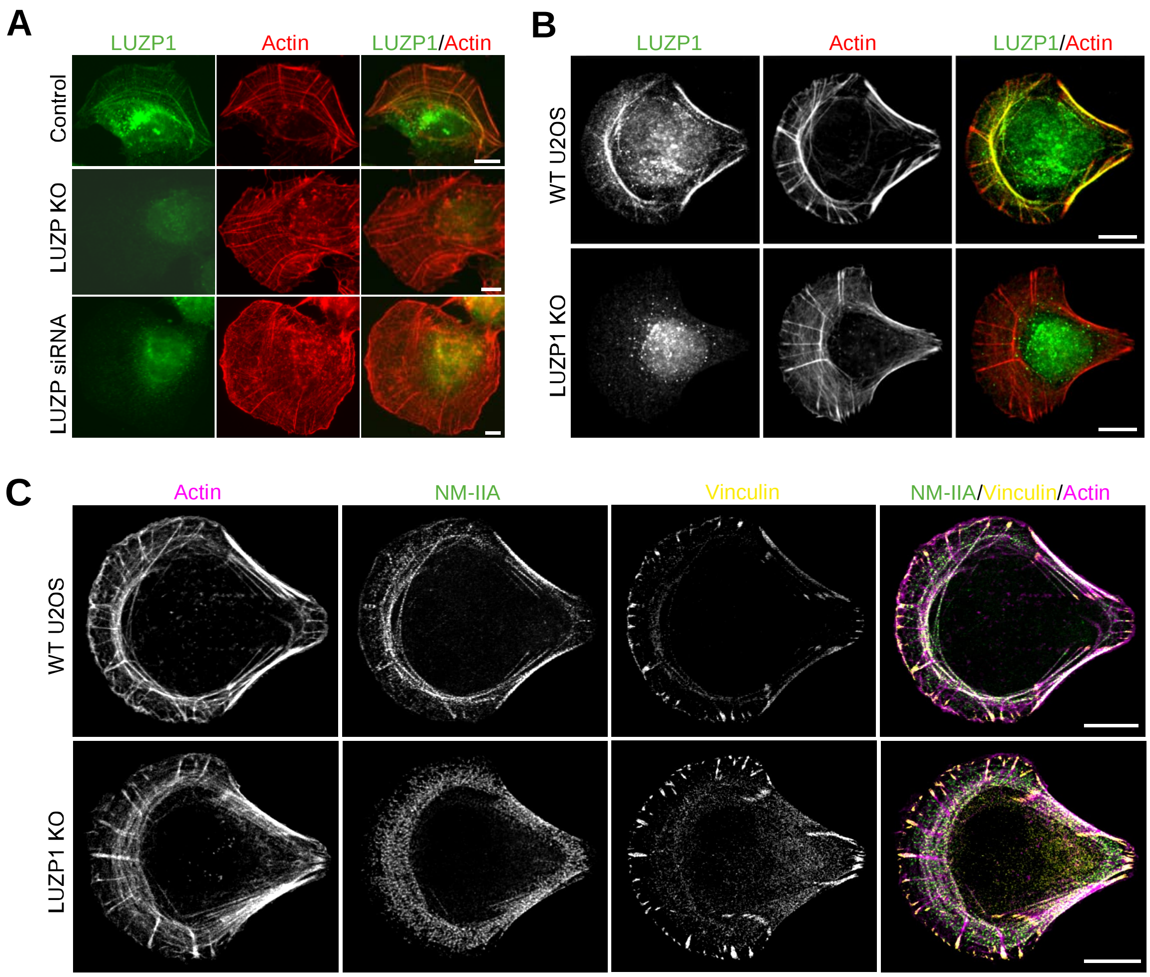

A
LUZP1
Actin
LUZP1/Actin
Control
LUZP KO
LUZP siRNA
B
LUZP1
Actin
LUZP1/Actin
WT U2OS
LUZP1 KO
C
Actin
NM-IIA
Vinculin
NM-IIA/Vinculin/Actin
WT U2OS
LUZP1 KO

### Figure S6

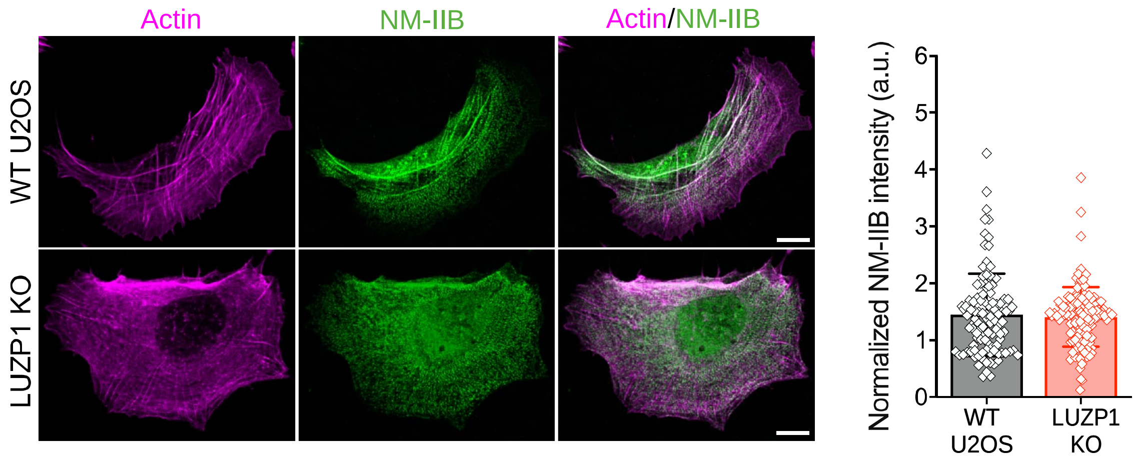
Actin
NM-IIB
Actin/NM-IIB
WT U2OS
LUZP1 KO
Normalized NM-IIB intensity (a.u.)
0
1
2
3
4
5
6
WT U2OS
LUZP1 KO
